# Granular Extracellular Matrix (gECM) Hydrogels Enable Distinct Composition and Mechanics Across Tissue Types for Translation

**DOI:** 10.64898/2026.05.06.723348

**Authors:** Juliet O. Heye, Shannon A. Blanco, Stephanie E. Schneider, Aseem M. Visal, Faith O. Olulana, Emily Y. Miller, Jeanne E. Barthold, Carson J. Bruns, Maxwell C. McCabe, Sean P. Maroney, Kirk C. Hansen, Corey P. Neu

## Abstract

Biomaterials-based tissue engineering aims to recapitulate native tissue architecture and function for both clinical repair and advanced *in vitro* models. While improvements in biomaterials have been made, including granular hydrogels and ECM-derived scaffolds, current biomaterials lack intentional design choices for effective translation, including regulatory considerations, practical extrusion delivery, and biomimetic characteristics. Here, we develop and characterize a library of granular ECM (gECM) biomaterials for five key tissues (cartilage, bone, skin, liver, and kidney), in which ECM particles are densely packed within a hyaluronic acid hydrogel. We optimize tissue processing methods that preserve proteomic content and structure while also aligning with scale-up manufacturing and regulatory guidelines. We show that gECM hydrogels can be molded, extruded, and 3D-printed while retaining their shape, and they stabilize at physiological temperature and pH. Lastly, we demonstrate that bulk gECM mechanics are driven by tissue type, and gECM hydrogels support viability, proliferation, and tissue-specific cellular activity. Together, these findings establish gECM hydrogels as a translational and biomimetic platform for clinical tissue repair and complex *in vitro* models.

## INTRODUCTION

Biomaterials-based tissue engineering aims to address a range of tissue disorders by replicating native tissue architecture and function, for both clinical and *in vitro* use. Cartilage, bone, skin, liver, and kidney make up about 80% of market applications, covering a broad range of disease systems [1]. For example, cartilage and bone are commonly affected by orthopedic conditions, including osteoarthritis [2]. Skin plays a major role in wound healing and plastic surgery [3], while the kidney and liver are critical for organ transplantation and drug metabolism [4]. One goal is to engineer tissues that clinically replace or repair damaged areas, restoring function in patients. In clinical applications, translational constraints must be considered, including processing requirements, cytocompatibility, and manufacturing time [5,6]. Engineered tissues can also be used to create 3D models that are essential for understanding disease pathogenesis and validating emerging therapeutics. The FDA Modernization Act 2.0 was passed in 2022, allowing for alternatives to animal testing for drug and biological product applications, including cell-based assays and microphysiological systems [7]. The recent regulatory shift creates an immediate need for reliable models for therapeutics testing [8]. As the demand for new tissue engineering solutions increases, there is a corresponding need to develop biomaterials that mimic native tissue architecture, can be easily delivered through injection or bioprinting, and are de-risked for translation.

Granular hydrogels are a relatively new class of materials following a bottom-up approach to biomaterial design in which building blocks, or granular particles, are packed densely together. Most commonly, granular hydrogels consist of densely packed microgels, or pre-polymerized hydrogel particles. These microgels are often formed through emulsion polymerization of synthetic or biopolymers and biological factors that are crosslinked together [9–11]. Some studies have formed microgels out of ECM-derived hydrogels [12,13], but this soluble form often requires the addition of synthetic crosslinkers to polymerize and lacks a biomimetic microstructure. Granular biomaterials demonstrate an ability to hold their shape upon injection, inter-particle porosity, and multiscale modularity–qualities that are difficult to achieve in a more traditional bulk hydrogel [14]. However, the micro-structure and composition within each microgel often lack the complex organization of components seen in native ECM. An alternative methodology is to make biomaterials from existing human or animal tissue with a top-down approach [2,15–18]. The tissue is typically decellularized to remove foreign DNA, and it is commonly enzymatically digested into soluble form [19,20]. While the complex array of ECM components is present, solid decellularized biomaterials lack the tunable mechanical properties of granular and engineered alternatives, whereas soluble versions lack the inherent structural architecture and mechanical integrity found in native ECM.

Combining these strategies, ECM can be mechanically processed into ECM particles and packed into a hydrogel base to form granular ECM (gECM) hydrogels with enhanced extrudability and structural biomimicry [21–24]. ECM particles have been incorporated into biomaterial scaffolds for a variety of tissue applications to preserve native tissue-specific biochemical and mechanical signals [25–31]. Notably, gECM hydrogels have demonstrated superior mechanical performance. For example, a liver-derived particulate ECM bioink exhibited more than a ninefold increase in elastic modulus compared to its solubilized counterpart [24]. ECM particles represent a promising direction for developing biomimetic, extrudable biomaterials that could enhance clinical and *in vitro* outcomes. Our team has previously developed a method to decellularize porcine cartilage with a detergent (2% SDS) and pulverize the tissue to form ECM particles [2,18]. We optimized the packing density of ECM particles into a hyaluronic acid base, establishing a formulation in which particles are packed beyond a percolation threshold to improve mechanical properties [18]. Importantly, we found that thiolated hyaluronic acid (tHA) directly crosslinks to ECM particles via disulfide bonding under physiological conditions and without synthetic crosslinkers [2]. We validated these materials with cell studies demonstrating that bovine chondrocytes infiltrate cartilage gECM hydrogels within 48 hours and express genes consistent with native cartilage repair after 14-day culture [18].

In this study, we expand our gECM hydrogel library by developing formulations for five key tissues: cartilage, bone, skin, liver, and kidney. We incorporate translational requirements for biomaterial processing, including the addition of viral inactivation, as required by the FDA for nonhuman tissues [5], and removal of detergents in our decellularization, which often leave residuals leading to failed cytotoxicity tests required to translate new materials for clinical use [32]. We also reduce our processing time to fit in a typical manufacturing window (<8hr shift), and separate animals to track donor variability and optimize biomaterial reproducibility. We characterize structural, biophysical, and mechanical properties of our gECM hydrogel library, and further demonstrate cellular responses for a subset of connective tissues (cartilage and skin). Ultimately, our work aims to create a versatile platform for tissue engineering that can support a range of applications, from disease modeling and drug testing to clinical tissue repair.

## RESULTS

### Detergent-free decellularization successfully reduces DNA content

We established a decellularization protocol for cartilage, bone, skin, liver, and kidney using an acid-base viral inactivation treatment [33] that is completed within 8 hours (**Figure 1A**). Confocal microscopy shows a reduction of DAPI-stained DNA following decellularization, suggesting the removal of cellular material (**Figure 1B**, **Supplementary Figure 1**). Quantification of DNA content (**Figure 1B**) confirmed a reduction of >98.5% between native and decellularized tissues, with decellularized tissue below 50 ng dsDNA per mg dry tissue [34].

**Figure 1.**
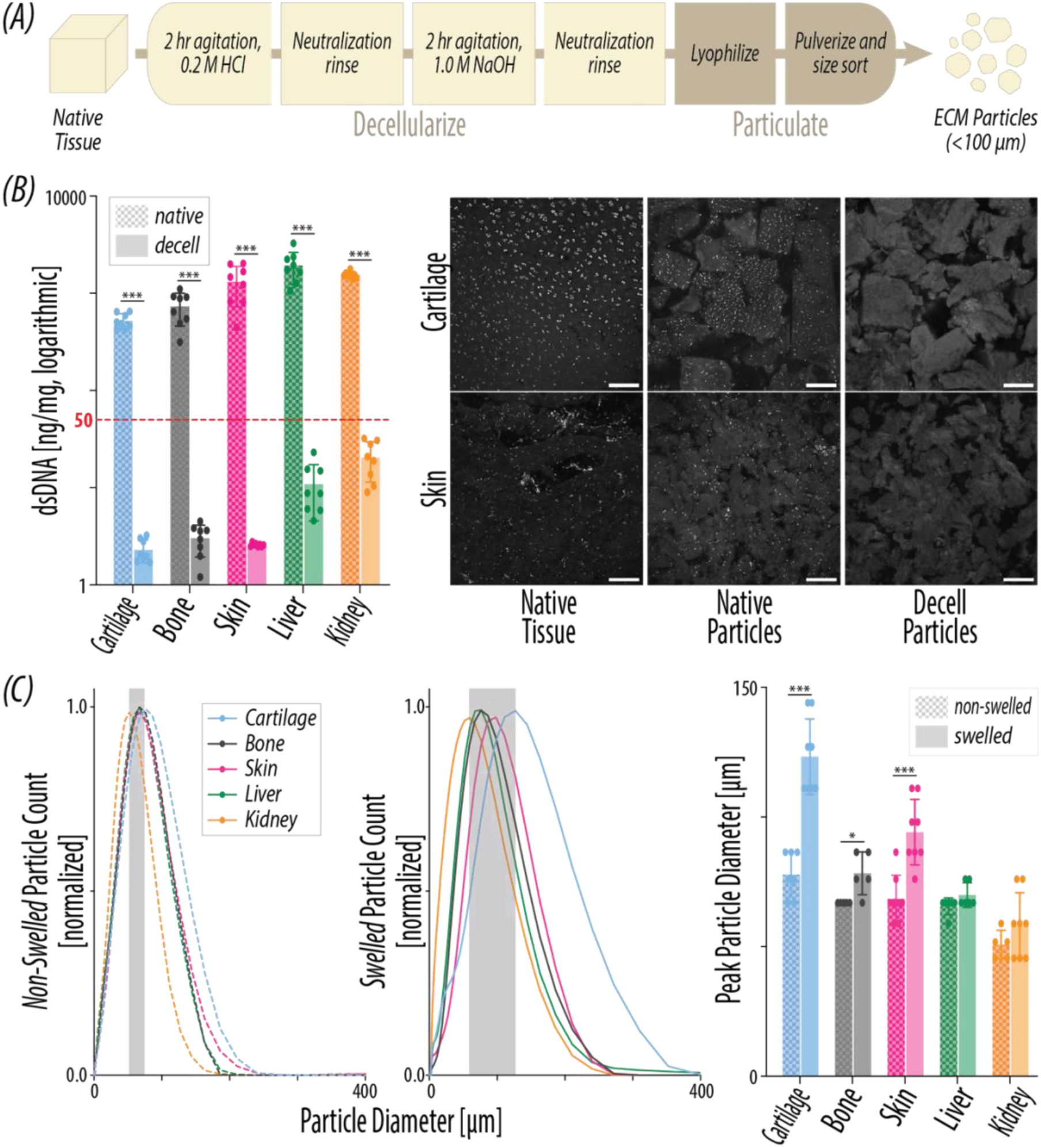
Tissue processing yields decellularized and size-sorted ECM particles. (A) Tissue is decellularized in under 8 hours and pulverized into <100µm gECM particles. (B) Double-stranded DNA (N = 8) content before (native) and after (decell) decellularization. Representative confocal images of cartilage and skin tissue showing DAPI-stained nuclei (white) in native whole tissue, native ECM particles, and decell ECM particles. Scale bars = 200 µm. (C) Size distribution of gECM particles in non-swelled and swelled states (N = 5-8). Peak particle size of gECM particles in non-swelled and swelled states (N = 4-8). Error bars = standard deviation. For all plots, *p<0.05, **p<0.01, ***p<0.001.

Notably, we did not use detergents in our decellularization protocol to mitigate the risk of adverse biocompatibility outcomes from residual detergent [32]. We compared cartilage processed with our <8-hour viral inactivation protocol to cartilage processed with an overnight viral inactivation and detergent method (**Supplementary Figure 2**). DAPI staining revealed high pixel saturation on particles treated with SDS, suggesting nonspecific binding of the DAPI stain to residual detergent. Detergent-treated cartilage gECM exhibited higher compressive modulus and viscosity compared to viral inactivation-only treated cartilage gECM. Proteomics reveal that the longer processing strips away ECM components like proteoglycans, leaving behind a collagen-dense and stiffer structure.

### Manufacturing of ECM particle size is tightly controlled, but increases when hydrated

While ECM particles are size sorted in dry form, their downstream use is in clinical or *in vitro* aqueous environments. As such, we then characterized the size distribution of ECM particles before and after hydration (**Figure 1C**). Non-swelled peak particle sizes for all tissues fall under 100 µm and are tightly clustered, spanning 27.12 µm across tissues. This is in line with the 100 µm sieve used to sort lyophilized particles and suggests tight control during processing. Swelled cartilage, bone, and skin exhibited a significant increase in size compared to their non-swelled counterparts, resulting in a wider 65.11 µm span of peak particle sizes across tissue types. These results suggest that different tissue types have different swelling capabilities, with cartilage particles increasing in size by 1.58-fold, bone by 1.17-fold, skin by 1.38-fold, liver by 1.07-fold, and kidney by 1.15-fold. We observed the largest swelling in cartilage and skin particles, likely due to their higher proteoglycan content, which influences the particle swelling behavior [35–38]. Particle swelling is an important characteristic to understand, as it may impact practical attributes (e.g., extrusion resolution) and biomimetic performance (e.g., mechanics and cell interactions).

### Decellularization of ECM particles retain key matrisome components, including collagens

Proteomics analysis (**Figure 2**, **Supplementary Figure 3**) further confirms a reduction to <2% cellular protein content after decellularization. Liver and kidney demonstrate an especially notable reduction in cellular content from >55% of total LFQ signal to <2% after decellularization. Proteomic analysis of matrisome components revealed that collagens constitute the dominant protein class across all tissues, with decellularization resulting in a reduction of non-collagen matrisome proteins. Despite this shift, decellularized cartilage, bone, and skin retained comparable proportional distributions among the remaining non-collagen matrisome protein categories, indicating preservation of tissue-specific ECM composition. In contrast, decellularized liver and kidney exhibited an increase in the relative proportion of ECM glycoproteins compared to their native counterparts, driven primarily by a reduction in proteoglycans in both tissues and a decrease in ECM regulators in liver. Abundance of the top 10 native proteins by LFQ signal is well-maintained in decellularized cartilage, skin, and bone. In decellularized liver and kidney, abundance in the top 3 proteins is maintained, with some reduction in subsequent proteins. A PCA plot (**Supplementary Figure 3A**) shows that cartilage and bone have distinct overall proteomic profiles compared to skin, kidney, and liver, which are clustered together. These results align with the top 10 proteins for each tissue type, with skin, kidney, and liver showing the same top 3 proteins (COL1A1, COL1A2, COL3A1).

**Figure 2.**
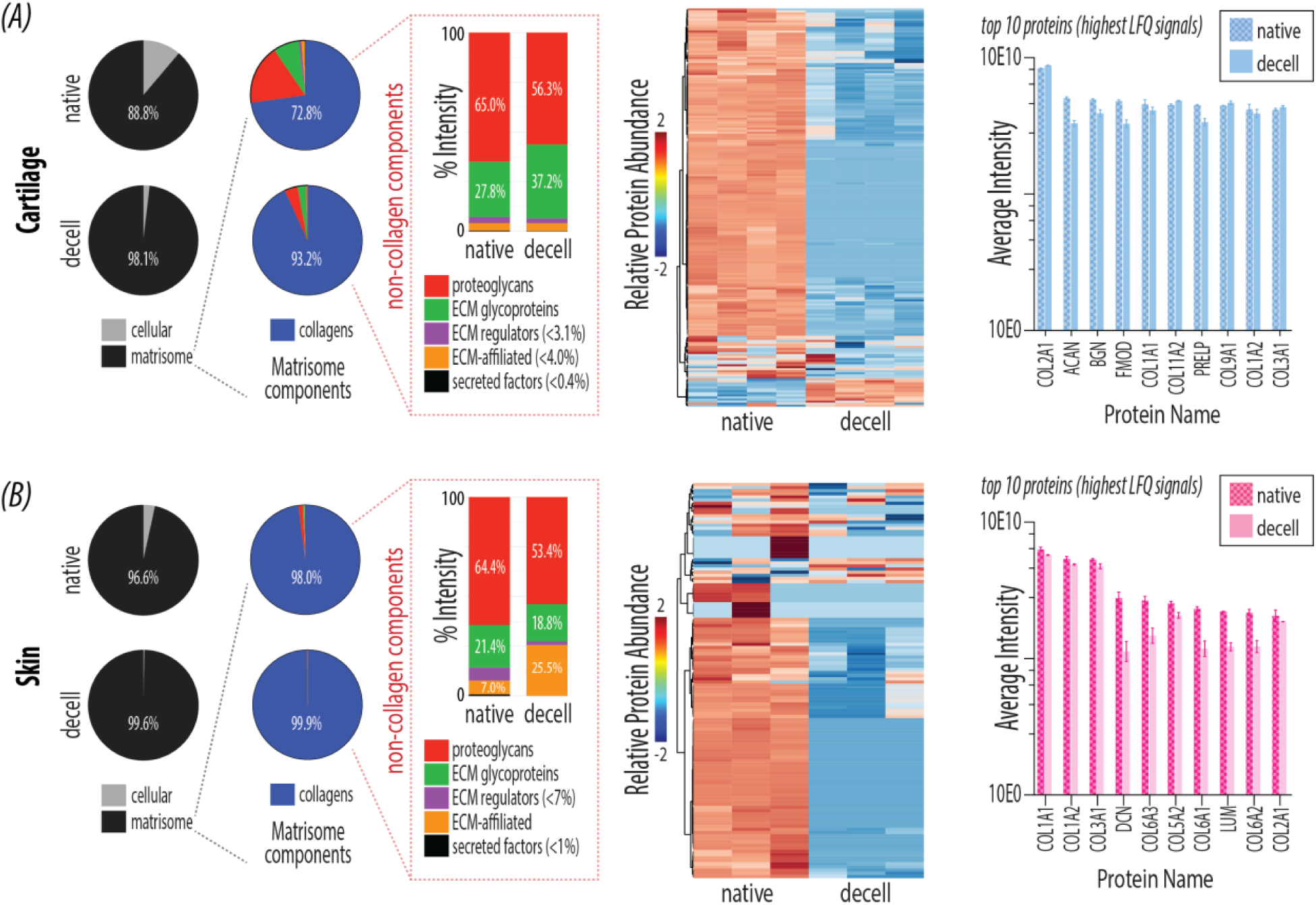
Decellularization of ECM particles retain key matrisome components, including collagens. (A) Cartilage (N = 4) and (B) Skin (N = 3) ECM particles proteomics compared before (native) and after (decell) decellularization. For each tissue, we present cellular vs. matrisomal protein content by LFQ signal. Remaining plots portray matrisome proteins only – matrisome subcategory distribution by LFQ signal, heat map with relative protein abundance, and top 10 protein abundance. Error bars = standard deviation.

### gECM hydrogels physically swell after formation, with dense particle packing

For subsequent analyses, particles and thiolated hyaluronic acid (tHA) are mixed in a 2-syringe system to make gECM hydrogels and polymerized in cylindrical molds to form constructs for testing (**Figure 3A**). All tissue gECM hydrogels maintain a pH of 7-7.5 after mixing and appear white in color (**Supplementary Figure 4**). We assessed swelling of polymerized gECM hydrogel constructs normalized to the constructs immediately after polymerization (0hr timepoint) (**Figure 3B**), to understand how the overall volume and mass of these materials change in an aqueous environment. Cartilage, bone, liver, and kidney gECM plateaued in volume after 1hr, with significant increases in volume only between 0hrs to 1hr. Skin gECM volume plateaued after 24hrs, with an additionally significant increase from 1hr to 24hrs. Cartilage, skin, and kidney gECM significantly decreased in volume between 1wk and the dry timepoint. When looking at swelling mass, cartilage and liver gECM plateaued after 1hr (significant increase 0-1hr), while bone and skin gECM plateaued after 24hrs (significant increase 1-24hrs), and kidney gECM plateaued after 48hrs (significant increase 1-48hrs). Dry mass was significantly lower for all gECM compared to all time points, which reveals the high water content of the gECM hydrogels. Overall, we see that the largest increase in volume and mass occurs in the first hour, with some gECM hydrogels, notably skin, continuing to swell beyond that time point. While tHA contributes to swelling behavior, variation is likely due to tissue-specific capacity for water uptake by the ECM particles. Notably, all gECM hydrogel constructs stayed intact throughout the swelling period, which suggests minimal risk of gECM detaching in a clinical or *in vitro* application. We additionally normalized swelling volume and mass to corresponding dry values and show raw measurements (**Supplementary Figure 5**).

**Figure 3.**
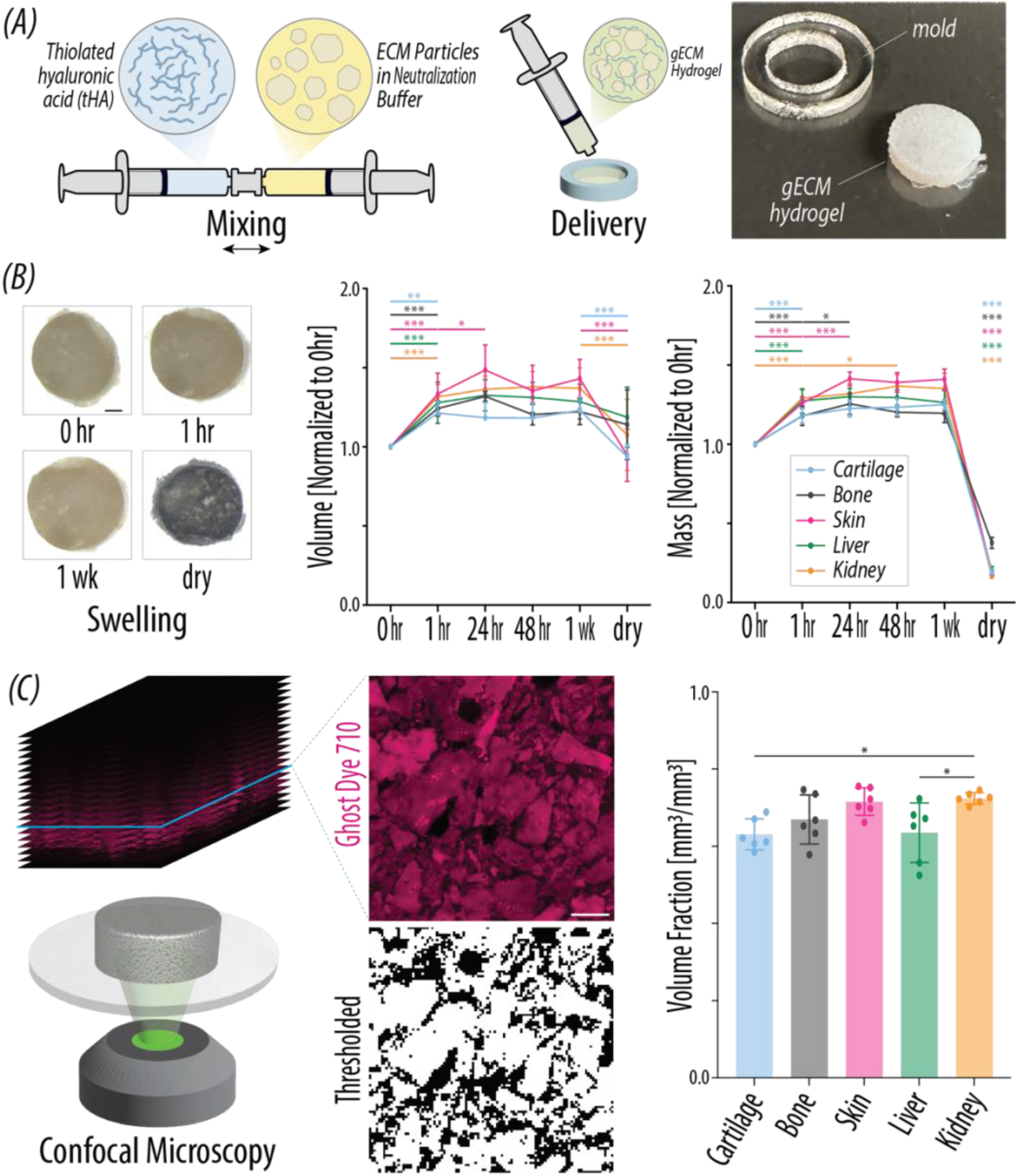
gECM hydrogels physically swell after formation, with dense particle packing. (A) gECM hydrogels are made by mixing thiolated hyaluronic acid (tHA) and ECM particles in neutralization buffer together using syringes. The mixed gECM hydrogel is extruded into a PDMS mold and polymerized to form cylindrical constructs for characterization. (B) Representative top-down images of gECM hydrogel samples during swelling, used to quantify surface area for volume calculations. Scale bar = 1 mm. Mass and volume of gECM hydrogels over the course of 1 week swelling in DPBS, normalized to initial volume immediately after mixing and polymerization (N = 4-6). (C) Representative confocal slices through optical depth of polymerized and swelled cartilage gECM hydrogel sample. Selected slices were thresholded in MATLAB, and area fraction of particles was averaged across slices to calculate volume fraction of particles to total volume (N = 6). Scale bar = 100 µm. For all plots, *p<0.05, **p<0.01, ***p<0.001.

We then quantified particle volume fraction for each tissue gECM (**Figure 3C**). Statistically, kidney gECM had a significantly higher volume fraction compared to cartilage and liver gECM. The high volume fraction for kidney gECM may be a result of small particle sizes (**Figure 1C**) that can more tightly pack together. Importantly, the volume fractions for bone, skin, liver, and kidney gECM exceeded that of cartilage gECM, which we use as a benchmark for geometric percolation [18] where particles have formed a network.

### gECM hydrogels are shear-thinning regardless of tissue type

Particle volume fraction also influences viscosity, which indicates extrudability. Viscosity of gECM hydrogels immediately after mixing showed shear-thinning behavior, with viscosity decreasing as shear rate increased (**Figure 4A**). No significant differences in viscosity curves were observed between the different tissue types (p=0.9508), suggesting that gECM tissue type does not drive viscosity. Shear-thinning behavior enables practical extrusion delivery like injection or 3D bioprinting, verified by successful extrusion of gECM hydrogels in a pneumatic 3D bioprinter (**Figure 4A**).

**Figure 4.**
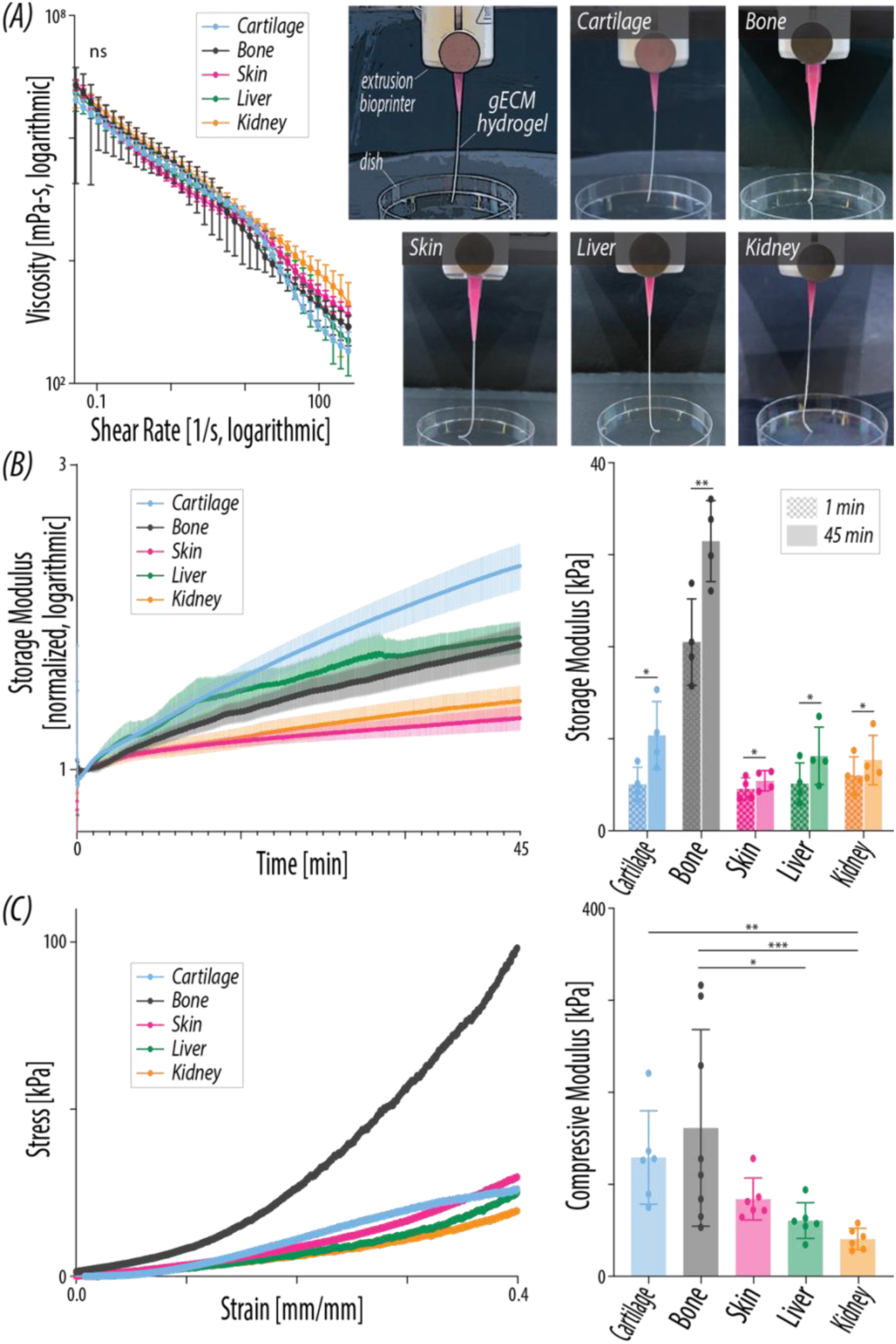
gECM hydrogels are shear-thinning but solid-like upon mixing, and stabilize over time with mechanics driven by tissue type. (A) gECM hydrogels demonstrate shear-thinning viscosity (N = 6). Representative photos of gECM hydrogels being extruded through a pneumatic 3D bioprinting nozzle. (B) Storage modulus of gECM hydrogels over 45 mins at 37°C, normalized to storage modulus at 1 min (N = 4). Error bars = SEM. Storage modulus (kPa) of gECM hydrogels at 1 min and 45 min at 37°C (N = 4). (C) Representative stress-strain curves of gECM hydrogels during bulk unconfined compression. Bulk compressive moduli of gECM hydrogels calculated between 20-30% strain (N = 6-8). Error bars = standard deviation. For all plots, *p<0.05, **p<0.01, ***p<0.001. ns = not significant.

### gECM hydrogels display solid-like rheological behavior even before polymerization and continue to stabilize in physiological conditions over time

We assessed rheological properties of the gECM hydrogels immediately after mixing components for 45 minutes at 37°C (**Figure 4B**), with oscillatory strain and frequency determined by preliminary amplitude and frequency sweeps [39] (**Supplementary Figure 6**). Loss modulus remains steady and below the storage modulus for all tissues across the entire 45-minute test (**Supplementary Figure 6**), suggesting a physically entangled particle network that makes the gECM behave like a solid even when uncured [40,41]. The gECM hydrogels hold their shape upon extrusion, which further verifies the observed solid-like rheological behavior. Storage modulus, normalized to the value at 1 minute [42] (**Figure 4B**), showed continuous stabilization over time, likely driven by thiol-crosslinking [2]. Storage modulus significantly increased between 1 and 45 mins for all gECM. Notably, cartilage exhibited the largest increase, potentially due to the quantity of thiol groups on the ECM particles available for crosslinking with tHA [2]. The relative ranking of storage moduli (bone > cartilage > skin/liver/kidney) aligns with the known relative stiffnesses of these tissues [43], indicating tissue particle stiffness may drive storage modulus. Tissue-specific particle size and shape could also modulate storage modulus under oscillatory strain [44].

### Tissue type drives bulk gECM stiffness

After polymerization, we measured bulk gECM stiffness. Representative stress-strain curves reveal a J-shape for all tissues except cartilage, which appears to be more S-shaped (**Figure 4C**). This may be due to a reorganization or slipping of particles at high strains in cartilage gECM, while the other tissues maintain a more stable network of particles. We calculated compressive modulus at 4 regions of strain (0-10%, 10-20%, 20-30%, and 30-40%) (**Supplementary Figure 7**). Overall, the compressive moduli increase with increasing strain. The relative ranking of compressive moduli among bone, skin, liver, and kidney (bone > skin > liver > kidney) was preserved across all regions of strain, but the relative ranking of cartilage varied depending on strain region, as expected based on the stress-strain curves. We highlighted the compressive moduli calculated at 20-30% strain, where the effect of percolation is exaggerated, but before the cartilage gECM appears to slip beyond 30% strain. Bone was significantly stiffer than liver and kidney, and cartilage was significantly stiffer than kidney. The relative ranking and differences in tissue compressive moduli support the hypothesis that tissue type drives bulk stiffness of gECM hydrogels [45], likely through geometrically percolated particles modulated by inherent ECM composition and stiffness [18]. We do see a relatively high standard deviation for bone compressive modulus, suggesting high donor variability with the bone gECM.

### gECM hydrogels support mechanical loading and cellular viability in a microphysiological model system

Two proof-of-concept microphysiologic chips (osteochondral and skin) with applied physiologically-relevant mechanics were developed to demonstrate potential applications for gECM hydrogels. The skin model (**Figure 5A**) consists of skin gECM flowed into the central chamber of an Emulate chip [46], with lateral vacuum chambers to apply stretch to the gECM. Deformation microscopy, an image-based technique used to calculate material deformation with and without load [47], was performed on confocal images before and after applied stretch, revealing tensile strain within the gECM.

**Figure 5.**
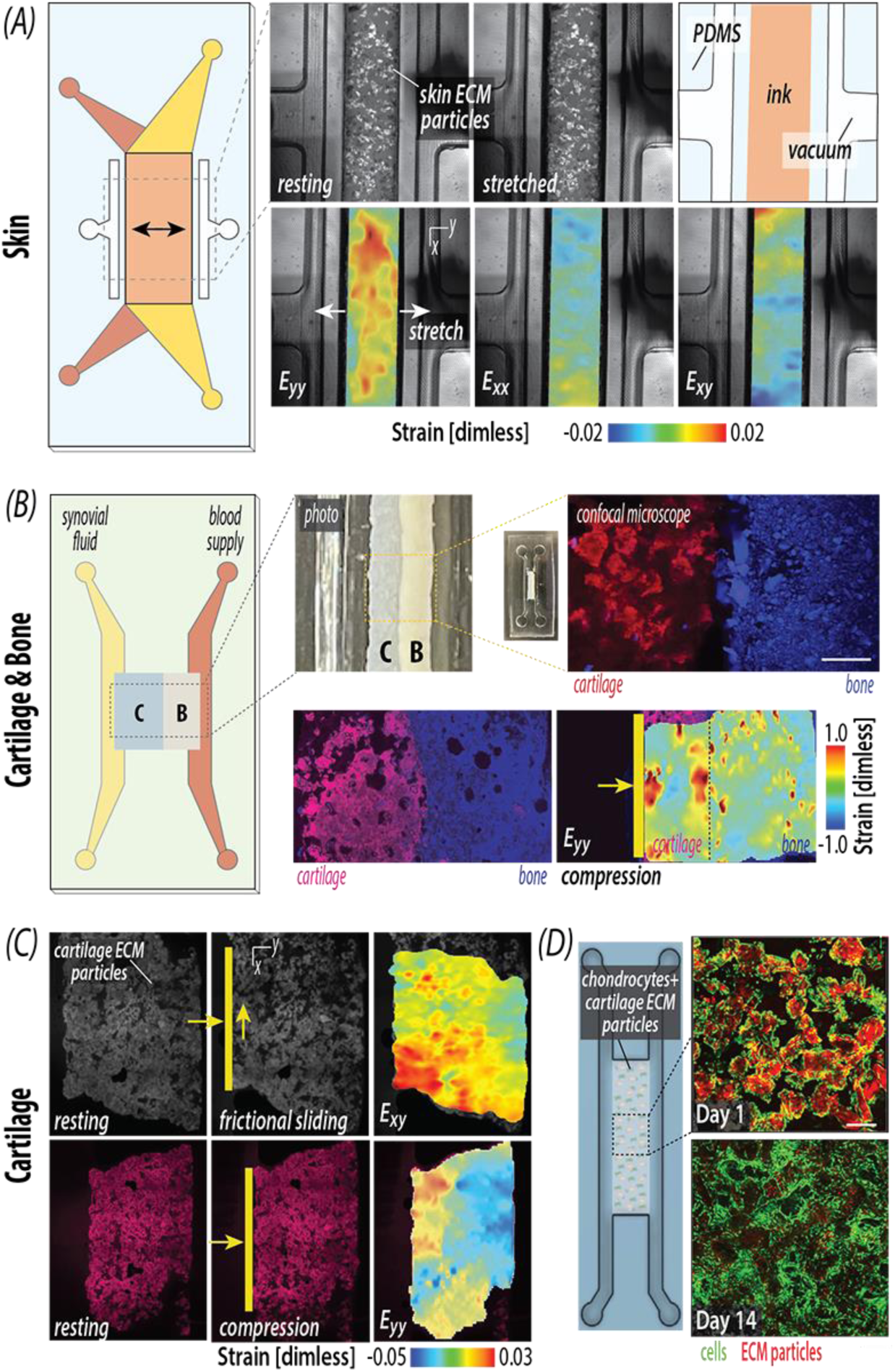
gECM hydrogels support mechanical loading and cellular viability in a microphysiological model system. (A) Skin gECM hydrogel was injected into a microphysiological Emulate chip, with lateral vacuum chambers to apply stretch. Deformation microscopy was performed on confocal images acquired before and after applied stretch to calculate strain maps. (B) Cartilage (left) and bone (right) gECM hydrogels were printed into a microphysiological osteochondral chip that was molded with PDMS. Scale bar = 10 mm. Confocal imaging of tissue-on-chip interface showing pre-stained cartilage (left, red) and bone (right, blue) gECM. Scale bar = 200 µm. Strain maps calculated by deformation microscopy of the cross-section of composite cartilage+bone gECM construct under compression, Eyy (cartilage red, bone blue). (C) Strain maps calculated by deformation microscopy of the cross-section of cartilage gECM under compression plus shear force (Exy), and compression only (Eyy). (D) Bovine chondrocytes cultured with cartilage gECM in a microphysiological chip, with confocal imaging at Day 1 and 14 of culture (green = calcein, red = ethidium homodimer). Scale bar = 200 µm.

The osteochondral chip (**Figure 5B**) was designed to contain side-by-side bone and cartilage layers, with individual fluid channels representing blood flow and synovial fluid, respectively. Bone and cartilage gECM were 3D-printed into the chip, and confocal microscopy revealed a distinct interface between pre-stained bone and cartilage gECM. We then visualized the mechanics of cartilage and bone gECM hydrogels by applying compression and shear force to polymerized constructs with an external microactuator. Cartilage-only gECM under compression, and under compression plus shear (**Figure 5C**) showed spatially-dependent mechanical signaling within the cartilage construct. Deformation microscopy of a composite cartilage and bone gECM construct (**Figure 5B**) showed higher strains in the y-direction in cartilage compared to bone gECM, as would be expected in a biological osteochondral system in which the cartilage deforms more than the bone. Mechanical stimulation, such as stretch in skin [48,49] and compression and shear within joints [50,51], is critical for recapitulating physiological systems. The spatial strain patterns observed under load in skin and osteochondral gECM hydrogels highlight their potential to model tissue for both clinical and *in vitro* applications.

Finally, we cultured primary bovine chondrocytes with cartilage gECM in our cartilage chip (**Figure 5D**). Over a 14-day period, these chondrocytes demonstrated viability, proliferation, and adherence to cartilage particles. These preliminary results support future use of our gECM hydrogels in cellularized on-chip *in vitro* models.

### gECM hydrogels maintain tissue-specific cellular gene expression *in vitro*

We cultured bovine chondrocytes encapsulated in cartilage gECM and murine dermal fibroblasts in skin gECM for 14 days. Confocal imaging showed high viability and proliferation from 3 to 14 days (**Figure 6A**). Cells retained their tissue-specific morphology, with chondrocytes appearing smaller and more rounded, while fibroblasts appeared spindle-shaped and spread out. Gene expression analysis at Day 3 and 14 (**Figure 6B**) revealed upregulation of key chondrogenic markers (SOX9, ACAN, COL2A1, PRG4) [18] in chondrocytes cultured in gECM compared to those cultured on tissue culture plastic (TCP) for only 3 days. COL1A2 is initially downregulated, indicating hyaline chondrogenic activity as opposed to fibrocartilaginous or hypertrophic activity [18]. A temporal decrease in SOX9, ACAN, COL2A1, and PRG4, paired with a temporal increase in COL1A2, suggests a potential trend towards de-differentiation, a common challenge with culturing chondrocytes [37,38,52,53]. However, our 3-day TCP condition represents a conservative control, as chondrocytes maintained on TCP for the full 14-day duration would be expected to undergo even greater de-differentiation [38]. As such, the upregulation of canonical chondrogenic genes in gECM supports its ability to delay de-differentiation compared to traditional 2D culture systems. Gene expression analysis of dermal fibroblasts in skin gECM revealed downregulation of *Acta2*, indicative of reduced myofibroblast activation [54], alongside upregulation of *Pdgfra* and *Thy1*, markers commonly associated with dermal fibroblast populations [55]. We did not see any notable differences in *Vim* compared to dermal fibroblasts cultured on TCP. *Col1a1* was also upregulated, suggesting active matrix synthesis in the absence of a fibrotic phenotype [54]. Together, the gene expression suggests that skin gECM supports a non-fibrotic, ECM-producing phenotype consistent with a healthy dermal fibroblast population.

**Figure 6.**
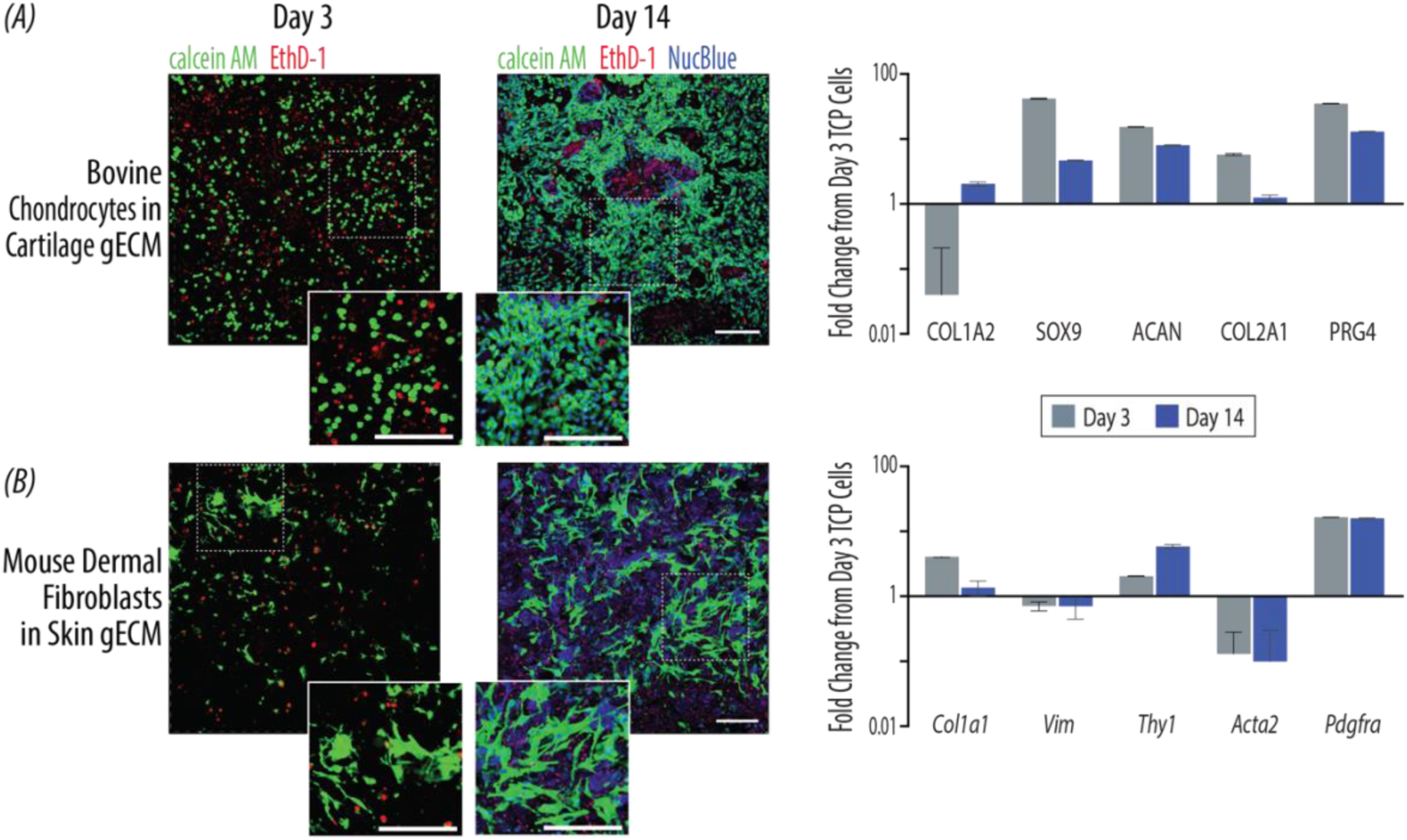
gECM hydrogels maintain tissue-specific cellular gene expression *in vitro.* (A) Bovine chondrocytes cultured on cartilage and (B) murine dermal fibroblasts cultured in skin gECM hydrogels (green = calcein, red = ethidium homodimer, blue = NucBlue). Scale bars = 200 µm. Quantitative PCR showing gene expression (N = 3), fold change to cells cultured on tissue culture plastic (TCP) for 3 days.

## DISCUSSION

In this study, we successfully implemented our gECM hydrogel technology for 5 porcine tissues (cartilage, bone, skin, liver, and kidney) to establish a platform for engineering disease models, drug testing, and clinical repairs. This study addresses recent advancements in ECM-derived biomaterials, in which ECM is particulated to maintain structural microarchitecture, and resulting particles are packed densely together to form granular ECM hydrogels. We placed strong emphasis on translational considerations in our gECM processing method to establish a protocol that is de-risked for manufacturing and regulatory pathways. We characterized practical gECM qualities, including extrudability, polymerization in physiological conditions, and swelling behavior, to enable easy use by researchers and clinicians. Finally, we demonstrate biomimetic gECM performance, showing that tissue-type modulates gECM mechanics. We also demonstrate the incorporation of gECM hydrogels into microphysiological models and provide evidence that cartilage and skin gECM support viability, proliferation, and tissue-specific gene expression of cells cultured on their respective matrix.

A core strength of the gECM platform lies in its alignment with translational and regulatory pathways. We established a detergent-free decellularization protocol that achieves consistent removal of cellular material across all tissues, with residual dsDNA levels below the 50 ng/mg threshold recommended for decellularized biomaterials [34]. By avoiding detergents, we reduce the risk of residual cytotoxicity [32] and streamline regulatory approval by limiting chemical inputs. The protocol also fits within an 8-hour window (i.e., a typical work day), making it feasible for large-scale manufacturing. Additionally, porcine tissue is abundant, inexpensive, and well-established as a clinical source material [56,57], facilitating production and widespread adoption. The gECM hydrogels presented here consist exclusively of natural, biologically-derived materials [2], improving biocompatibility and thereby reducing regulatory hurdles for clinical translation. We have also shown that the gECM platform can be adapted to diverse tissue types, offering a simple pipeline to produce gECM hydrogels for multiple tissue engineering applications. One limitation of our tissue processing is the need for a defatting step for porcine bone. Porcine bone had particularly high fatty bone marrow content, resulting in bone particles that demonstrated hydrophobic properties when exposed to tHA. As such, bone particles underwent an additional defatting process, which involved SDS. To mitigate the risk of residual detergent, future work should defat bone with an alternative process, such as hydrogen peroxide [58].

In parallel with translational de-risking, the gECM hydrogels exhibited practical characteristics relevant for clinical use and microphysiological model fabrication. Viscosity profiles confirmed that, regardless of tissue type, our gECM hydrogels are shear-thinning, fulfilling an important practical requirement for clinical or 3D modeling via extrusion delivery. We demonstrated successful 3D bioprinting of all 5 gECM tissues, including a proof-of-concept tissue-on-chip construct with bone and cartilage gECM. One limitation is that our biomaterials are limited in extrusion resolution by the particle sizes, but this could be mitigated in future preparations by sorting ECM particles to smaller (<100µm) diameters. We also qualitatively observed that the bone gECM hydrogel required higher forces to extrude. Because the viscosity profile was consistent across tissue types, we suspect this may be due to the heavy mineral content in the bone influencing how particles can move through the constrained geometry of the 1mL syringes used [59]. A custom-designed syringe may improve the smoothness of extrusion for bone gECM. Finally, we showed that polymerization occurs under physiological conditions without the need for external crosslinkers [2]. Rheological measurements demonstrated that even uncured gECM hydrogels have solid-like storage and loss modulus patterns, which support their ability to hold shape even in an uncured state. This allows researchers to mold, 3D print, or extrude the gECM hydrogels without concern about pre-polymerization stabilization, and enables surgeons to physically mold the gECM hydrogels to fit into irregular tissue defects. We demonstrated material stability by 3D printing bone and cartilage gECM directly into a PDMS microphysiological model. We also quantified volume and mass during swelling, enabling us to anticipate how gECM constructs will absorb liquid and change in volume in the body or in 3D culture models. This enables researchers and clinicians to make informed decisions on how to size a microphsyiological model, or whether to underfill a clinical defect and advise a patient on post-operative swelling. Importantly, our swelling study also revealed that the gECM constructs remain intact in an aqueous environment over time, mitigating the risk of gECM debris migration *in vivo* or *in vitro* [60].

The gECM hydrogels displayed biomimetic composition, mechanics, and cellular activity. While proteomic analysis revealed a reduction of non-collagen proteins, the relative proportion of protein types within the non-collagen protein category was preserved. This was further confirmed by maintenance of top 10 protein abundance after decellularization and distinct overall proteomic profiles, especially in cartilage and bone compared to skin, liver, and kidney. The differences in protein composition were reflected in mechanical behavior: bulk compressive modulus of the gECM hydrogels followed expected trends based on tissue origin [45], with bone gECM exhibiting the highest stiffness and kidney gECM the lowest. This observation supports a model in which gECM bulk stiffnesses arise from geometric percolation networks of particles [18], allowing us to capture mechanically distinct properties based on tissue type. Previously in our lab, we used the stiffness of cartilage gECM hydrogels to find a percolation threshold, defined by a specific volume fraction of particles to total gECM volume at which the particles form a network and drive overall mechanics [18]. We controlled particle volume fraction across tissue types, using the cartilage gECM geometric percolation as a benchmark for particle connectivity. It’s important to note that we have not measured and calculated a true percolation threshold for each tissue. Rather, we use cartilage as a benchmark. Different tissue particle shapes and sizes may influence the percolation threshold for specific tissues [44]. Regardless, we found that all tissue gECM hydrogels reached the cartilage percolation threshold and demonstrated expected tissue-specific differences, suggesting that a connective network has formed in the gECM hydrogels. It is important to acknowledge that gECM hydrogels, while microstructurally complex, display a homogenous bulk structure and thus do not inherently represent the complex macrostructural organization of native tissues. However, these gECM hydrogels can serve as building blocks to fabricate the macrostructure of (e.g., layered) tissue via 3D bioprinting or incorporation into on-chip systems. For example, we demonstrated assembly of our cartilage and bone gECM hydrogels into a microphysiological chip system with applied compression and load, mechanical features that create a more comprehensive representation of native tissue. Additionally, we showed that chondrocytes could be loaded onto this chip to establish a tissue-realistic model.

Finally, our gECM hydrogels promoted tissue-specific cellular behavior in two test cases – bovine chondrocytes encapsulated in cartilage gECM, and murine dermal fibroblasts encapsulated in skin gECM. The use of two different cell sources demonstrates that our porcine-derived gECM can promote tissue-specific behavior across a range of cell origins. Chondrocytes encapsulated in cartilage gECM for 14 days exhibited upregulation of key chondrogenic markers, including SOX9, ACAN, COL2A1, and PRG4 [18,38], compared to chondrocytes seeded in 2D on TCP for only 3 days. Although some markers declined over time, the overall upregulation relative to tissue culture plastic (TCP) underscores the benefit of gECM hydrogels for phenotype maintenance. Similarly, dermal fibroblasts cultured on skin gECM displayed increased expression of *Pdgfra* and *Col1a1*, with suppression of *Acta2*, suggesting a regenerative, non-fibrotic phenotype [54,55]. Our findings demonstrate that tissue-matched gECM hydrogels can promote tissue-specific morphology and gene expression.

gECM hydrogels offer characteristics important for downstream applications. Granular ECM hydrogels provide advantages compared to bulk polymer hydrogels, which have long been used in scaffold-based tissue engineering. Biopolymer or solubilized ECM hydrogels offer intrinsic cell-adhesive cues but can lack mechanical integrity, whereas synthetic hydrogels provide tunable mechanical and chemical properties but can lack biological familiarity. Granular hydrogels augment basic bulk hydrogels by introducing a multiscale structure that enables both tunable mechanics and composition. Compared to microgel-based granular hydrogels, gECM hydrogels directly preserve native extracellular matrix in their tissue microparticles, including microarchitecture and innate composition. This approach is also modular to a variety of tissue types, enabling use in many disease applications. We have demonstrated that gECM hydrogels can be flowed and 3D printed into microphysiological models, exhibit spatial strains under physiologically-relevant loading, and support cell viability and gene expression. With this foundation, multi-tissue microphysiological models could be created to understand disease pathology, like crosstalk between bone and cartilage in the onset of osteoarthritis, or to test the safety and efficacy of emerging drugs, like assessing toxicity in a kidney-based model. Clinically, gECM hydrogels have the potential to be noninvasively implanted, immediately withstand physiological loading, and guide cells to lay down new matrix over time. Potential disease applications include replacing cartilage in patients with injury or osteoarthritis and assisting soft tissue reconstruction in wound healing.

By leveraging the unique properties of tissue-derived ECM particles, we have developed a modular gECM hydrogel platform that can be tailored for specific clinical and research needs. To the best of our knowledge, we are the first to establish a granular ECM hydrogel in which the granules are made of particulated tissue ECM [2,17,18], instead of microgels. We have shown that the gECM technology is a versatile approach that can be widely applied to a variety of tissue types and that is well-positioned for clinical translation. Our findings motivate us to develop gECM hydrogels with human tissues in the future and to characterize their potential in guiding differentiation of human mesenchymal stromal cells (MSC). Future work with these gECM hydrogel can be used to engineer a scalable multi-component joint-on-chip system with applied load. In conclusion, our gECM hydrogels demonstrate translational, practical, and biomimetic characteristics that support their development as a promising platform for engineering new tissue.

## MATERIALS AND METHODS

### Tissue Particle Processing

#### Tissue sourcing and dissection

Porcine tissues were sourced from a local abattoir from sows ∼2 years old and separated by animal. Articular cartilage was dissected from the femoral condyles and trochlear groove of the stifle and cut into ∼4mm^3^ pieces. Trabecular bone was dissected using a chisel and hammer from the epiphysis and metaphysis of the femur in the stifle, and the epiphyseal line and cortical bone were removed. Skin was sourced from porcine underbelly, and subcutaneous fat was removed. Whole liver and kidney cortex was cut into small pieces. Bone, liver, and kidney pieces were orbitally agitated in DPBS overnight to rinse out blood. The bone, skin, liver, and kidney were then pulverized in a stainless-steel milling jar (Tissue Lyser III, Qiagen) and frozen until decellularization.

#### Decellularization

Tissues were decellularized using a viral inactivation method [33]. Tissues were orbitally agitated at RT in 0.2M HCl for 2hrs, then 1.0M NaOH for 2hrs, and rinsed in water/DPBS until neutral (pH=7.5). Prior to decellularization, skin underwent a 30min orbital agitation in 70% ethanol and 4×5min centrifugations in DPBS to remove fat. Additionally, bone underwent a de-fat process after decellularization, including a 3hr orbital agitation in 2% SDS, 8hr orbital agitation in ethanol (50%, 75%, 95%, 100% for 2hrs each) [58], and overnight orbital agitation in DPBS. Decellularized tissue was flash frozen, lyophilized, pulverized in a stainless-steel milling jar (Tissue Lyser III, Qiagen), and size sorted to <100 µm diameter using a sieve.

### Tissue Particle Characterization

*DNA content:* DNA content was quantified in lyophilized and pulverized native and decellularized ECM particles. Double-stranded DNA (dsDNA) content was extracted from processed cartilage and bone tissue particles (Zymo DNA Extraction Kit) and quantified using a PicoGreen assay (ThermoFisher Invitrogen). DNA content was visualized via confocal imaging (Nikon A1R, 10x objective, NA=0.45) of DAPI (ThermoFisher Invitrogen) stained tissues.

#### Particle Size and Swelling

Particle size distribution of gECM particles suspended in 100% ethanol (non-swelled) or DI water (swelled) was measured in a laser diffraction particle size analyzer (Mastersizer 3000). Mie theory was applied to calculate volume-based particle size distributions. Measurements were considered valid when residual error was <1%. The Beer-Lambert law was used to monitor optical obscuration and confirm appropriate sample concentration. Tissue-specific refractive indices were used (cartilage: 1.40) [61], bone: 1.55 [62], skin: 1.41 [63], liver: 1.37 [64], and kidney: 1.51 [65]), while the absorptive index (0.1) and density (1.00 g/cm^3^) were held constant across tissues [64,66–69].

#### Proteomics sample preparation

To accommodate different tissue types, native and decellularized bone, skin, liver, and kidney samples were processed using a sequential three-fraction extraction to isolate cellular, soluble extracellular matrix (sECM), and insoluble ECM (iECM) components, while native and decellularized cartilage samples were subjected directly to the hydroxylamine extraction step, generating only an iECM fraction. All subsequent extraction buffers were applied at a ratio of 200 μL/mg of the starting tissue dry weight. For three-fraction extraction, 3 mg of lyophilized material from each bone, skin, liver, and kidney sample was combined with 100 mg of 1 mm glass beads (Next Advance #GB10) and homogenized in decellularization buffer (50 mM Tris-HCl (pH 7.4), 0.25% CHAPS, 25 mM EDTA, 3 M NaCl, 1X Halt™ Protease Inhibitor Cocktail (Thermo Scientific #78438)) at power 8 for 3 min (Bullet Blender, Model BBX24, Next Advance, Inc.). The homogenate was vortexed (power 8) for 20 min at 4°C, spun at 18,000 x g (4°C) for 15 min, and the supernatant was collected. This decellularization process was repeated for a total of 3 washes, homogenizing and vortexing samples before each collection, and all washes were pooled to generate the cellular fraction. The remaining pellets were then homogenized in 6 M guanidine hydrochloride (Gnd-HCl), 100 mM ammonium bicarbonate (ABC) at power 8 for 1 min and vortexed (power 5) at room temperature overnight. This homogenate was spun at 18,000 x g (4°C) for 15 min, and the supernatant was collected as the sECM fraction. To extract the final iECM fraction, the remaining pellets from the bone, skin, liver, and kidney samples, as well as the independent lyophilized cartilage samples (0.5–2 mg), were treated with freshly prepared hydroxylamine buffer (1 M NH_2_OH−HCl, 4.5 M Gnd−HCl, 0.2 M K_2_CO_3_, pH adjusted to 9.0 with NaOH). The previously processed pellets were homogenized at power 8 for 1 min, while the intact cartilage samples were homogenized at power 8 for 3 min. All samples were subsequently incubated at 45°C with shaking (1000 rpm) for 4 h. Following incubation, the samples were spun for 15 min at 18,000 x g, and the supernatant was removed and stored as the iECM fraction. All collected fractions were immediately frozen and stored at −80°C until further proteolytic digestion. Extracted proteins were subjected to enzymatic digestion overnight (16h) at 37°C with trypsin (1:100 enzyme to protein ratio) using a filter aided sample preparation (FASP) approach as previously described [70] and desalted during Evotip loading.

#### LC-MS/MS analysis

Digested peptides (200 ng) were loaded onto individual Evotips following the manufacturer‘s protocol and separated on an Evosep One chromatography system (Evosep, Odense, Denmark) using a Pepsep column, (150 µm inter diameter, 15 cm) packed with ReproSil C18 1.9 µm, 120Å resin. Samples were analyzed using the instrument default “30 samples per day” LC gradient. The system was coupled to the timsTOF Pro mass spectrometer (Bruker Daltonics, Bremen, Germany) via the nano-electrospray ion source (Captive Spray, Bruker Daltonics). The mass spectrometer was operated in PASEF mode. The ramp time was set to 100 ms and 10 PASEF MS/MS scans per topN acquisition cycle were acquired. MS and MS/MS spectra were recorded from m/z 100 to 1700. The ion mobility was scanned from 0.7 to 1.50 Vs/cm^2^. Precursors for data-dependent acquisition were isolated within ± 1 Th and fragmented with an ion mobility-dependent collision energy, which was linearly increased from 20 to 59 eV in positive mode. Low-abundance precursor ions with an intensity above a threshold of 500 counts but below a target value of 20000 counts were repeatedly scheduled and otherwise dynamically excluded for 0.4 min.

#### Global proteomic data analysis

Data was searched using MSFragger v4.3 via FragPipe v23.1 [71]. Precursor tolerance was set to ±15 ppm and fragment tolerance was set to ±25 ppm. Data was searched against UniProt restricted to *Sus scrofa* with added common contaminant sequences [72] (46,127 total sequences, downloaded 9/25/25). Enzyme cleavage was set to semi-specific trypsin for all samples. Fixed modifications were set as carbamidomethyl (C). Variable modifications were set as oxidation (M), oxidation (P) (hydroxyproline), deamidation (NQ), Gln->pyro-Glu (N-term Q), and acetyl (protein N-terminus). Label free quantification was performed using IonQuant v1.11.11 with match-between-runs enabled and default parameters. Cellular, sECM, and iECM fractions were searched separately and merged after database searching. Results were filtered to 1% FDR at the peptide and protein level. Matrisome protein annotations were derived from MatrisomeDB [73] (annotation levels 1 and 2), in addition to in-house generated annotations for matrisome subcategories (annotation levels 3-5).

### gECM hydrogel Formulation

#### Hyaluronic Acid Functionalization

Glucoronate carboxyl groups on hyaluronic acid (Lifecore Biomedical, HA-100K-5) were replaced with thiol groups following previously established protocols to produce thiolated hyaluronic acid (tHA) [18], with purification via tangential flow filtration. The substitution rate was confirmed to be 18%–23% (thiolated mmols/unthiolated mmols) using a standard Ellman’s assay (Ellman’s solution, ThermoFisher).

*gECM hydrogel Mixing:* The gECM hydrogel formulations consisted of 10 mg/mL tHA packed with 0.2 g/mL (cartilage, skin, liver, kidney) [18] or 0.56 g/mL (bone) lyophilized particles. One syringe containing ECM particles in a neutralization buffer (DPBS + 1M NaOH) was mixed via luer lock connector to a second syringe containing an equal volume of 20 mg/mL tHA. For subsequent characterization of polymerized samples, gECM hydrogels were molded into cylindrical constructs (1.5 mm height x 5 mm diameter) and incubated for 45 min at 37°C.

### gECM hydrogel Biophysical Characterization

#### Swelling

Polymerized gECM hydrogel constructs were incubated in DPBS at room temperature for 1 week and then lyophilized. Mass and volume immediately after polymerization (0 hr), at 4 timepoints during incubation in DPBS (1 hr, 24 hrs, 48 hrs, 1 wk), and after lyophilization. Volume was obtained by measuring height with calipers and calculating surface area from thresholded (Image J) stereoscopic images (Leica).

#### Particle Volume Fraction

ECM particles in polymerized constructs were stained with Ghost Dye 710 (Cytek Biosciences) and imaged via confocal microscopy (Nikon A1R, 20x objective, NA=0.75). We obtained three-dimensional 5µm-step z-stack images at 3 different locations per sample. Using a custom MATLAB code, we selected, thresholded, and averaged the area over 15µm depth to calculate particle volume to total construct volume.

### gECM hydrogel Mechanical Characterization

#### Rheology

Within 5 minutes of mixing the gECM hydrogels, a room temperature rheological shear sweep (0.1 to 250 s^-1^) measuring viscosity was performed (MCR 702, Anton Paar). Separately, we performed a 45-minute time sweep measuring storage (G’) and loss (G’’) modulus at 37°C, with constant oscillation at 1% shear strain and 1 rad/s frequency (MCR 702, Anton Paar). Parameters were determined by identifying linear trends in preliminary amplitude (0.01-100% strain, 1 rad/s frequency) and frequency (0.01-100 rad/s, 1% strain) sweeps for gECM hydrogels in both uncured (immediately after mixing) and cured (after 45mins at 37°C) states. Rheological tests were performed with 8mm parallel plates and a 1.5mm gap height, which is >10x the peak particle diameter for all tissue types [39].

#### Extrusion Bioprinting

All tissue gECM hydrogels were 3D printed on a multi-nozzle pneumatic extrusion printer (BioX, Cell Ink Life Sciences) with 20 G nozzles, 40-80 kPa pressure, and a rate of 12 mm/s.

#### Bulk Compression

Polymerized gECM hydrogel constructs were compressed (unconfined) to 40% strain at a quasi-static 0.1%/s strain rate (MCR 702, Anton Paar). Compressive modulus was calculated between 0-10%, 10-20%, 20-30%, and 30-40% strain [18].

### Microphysiological Models Demonstration

#### Tissue-on-chip

Pre-stained skin gECM was flowed into an Emulate chip. Pre-stained cartilage (Ghost Dye 710) and bone (DAPI) gECM were 3D-printed on a multi-nozzle pneumatic extrusion printer (BioX, Cell Ink Life Sciences) into a molded Sylgard 184 polydimethylsiloxane (PDMS) tissue-on-chip model and imaged using confocal microscopy (Nikon A1R, 10x objective, NA=0.45).

#### Deformation Microscopy

The skin gECM was imaged via confocal microscopy (Nikon A1R, 10x objective, NA=0.45) before and after stretch that was applied via lateral vacuum chambers in the Emulate chip [46]. The cross section of a cartilage-only gECM construct, and stacked cartilage-bone gECM construct was stained and imaged via confocal microscopy (Nikon A1R, 10x objective, NA=0.45) during compression and shear with a micro-actuator. Deformation microscopy was used to compute construct strains using deformed and undeformed images [47].

#### Chondrocytes-on-chip

Primary bovine chondrocytes were sourced from juvenile bovine knee joints [37,38] and flowed into a microphysiological PDMS chip with cartilage gECM. After 45 mins at 37°C, media (Dulbecco’s Modified Eagle Medium: DMEM-F12 supplemented with 10% FBS, 0.1% bovine serum albumin, 100 units/mL penicillin, 100 µg/mL strep, and 50 µg/mL ascorbate-2-phosphate) was flowed through and replaced every 3 days during the culture period. We assessed viability (Calcein AM, Ethidium Homodimer-1; ThermoFisher Invitrogen) via confocal microscopy (Nikon A1R, 10x objective, NA=0.45) at Day 3 and 14.

### Cell Seeding

Primary bovine chondrocytes were sourced from juvenile bovine knee joints [37,38] and immediately seeded into 100 µL cartilage gECM hydrogels (7.5x10^6^ cells/mL). Primary murine dermal fibroblasts were sourced from the skin of embryonic mice and expanded to P3 before seeding into 100 µL skin gECM hydrogels (5x10^6^ cells/mL). Dry gECM particles were placed into a PDMS ring, cells suspended in 10 mg/mL tHA were pipetted into the ring, and constructs were incubated for 45 mins at 37°C before adding media (Chondrocytes: Dulbecco’s Modified Eagle DMEM-F12 supplemented with 10% FBS, 0.1% bovine serum albumin, 100 units/mL penicillin, 100 ug/mL strep, and 50 ug/mL ascorbate-2-phosphate; Dermal fibroblasts: Dulbecco’s Modified Eagle Medium DMEM supplemented with 10% FBS, 100 units/mL penicillin, and 100 µg/mL strep). We cultured the cell-laden gECM constructs for 14 days, changing medium every 3 days. On day 3 and day 14, we assessed cell viability (Calcein AM, Ethidium Homodimer-1, NucBlue; ThermoFisher Invitrogen) via confocal microscopy (Nikon A1R, 10x objective, NA=0.45) and gene expression through quantitative real-time PCR (CFX96 Touch). Samples were lysed in Qiazol and RNA was isolated using a Direct-zol RNA Miniprep Kit (Zymo). RNA was reverse transcribed into cDNA using iScriptTM (BioRad). Real-time quantitative PCR (RT-qPCR) was performed with SsoAdvanced^TM^ Universal SYBR® (BioRad) in a CFX96 thermocycler (BioRad). We targeted known genes to assess chondrogenic phenotype (SOX9, COL1A2, COL2A1, ACAN, and PRG4) and dermal fibroblast phenotype (*Col1a1, Vim, Thy1, Acta2, Pdgfra*). All measurements were normalized to the housekeeping gene (GAPDH), and fold changes were measured from gene expression of cells cultured on tissue culture plastic until confluent (∼3 days).

## Statistical Analysis

Mixed-model analyses of variance (ANOVAs) were performed on linear mixed models with tissue type and treatment (e.g. native vs decell, swelled vs non-swelled, etc) as co factors, animal donor as a random effect, and the resulting measurement variable as the response (DNA, particle size, swelling volume, swelling mass, volume fraction, viscosity, storage modulus, compressive modulus). All residuals were checked, and if a non-normal distribution was identified, the data was transformed for statistical analysis. Post hoc multiple comparisons were performed using Tukey’s honestly significant difference (HSD) correction. Significance was defined as p<0.05.

## Acknowledgments

The authors acknowledge Dahlia Ortiz for technical support on tissue decellularization and processing.

## Author contributions

Conceptualization: JOH, JEB, CPN

Methodology: JOH, SAB, SES, AV, FOO, EYM, JEB, CB, MCM, SPM, KCH, CPN

Visualization: CPN, JOH

Funding acquisition: CPN, JOH

Supervision: CPN, SES

Writing – original draft: JOH

Writing – review & editing: All authors

---

**Supplemental Figure 1.**
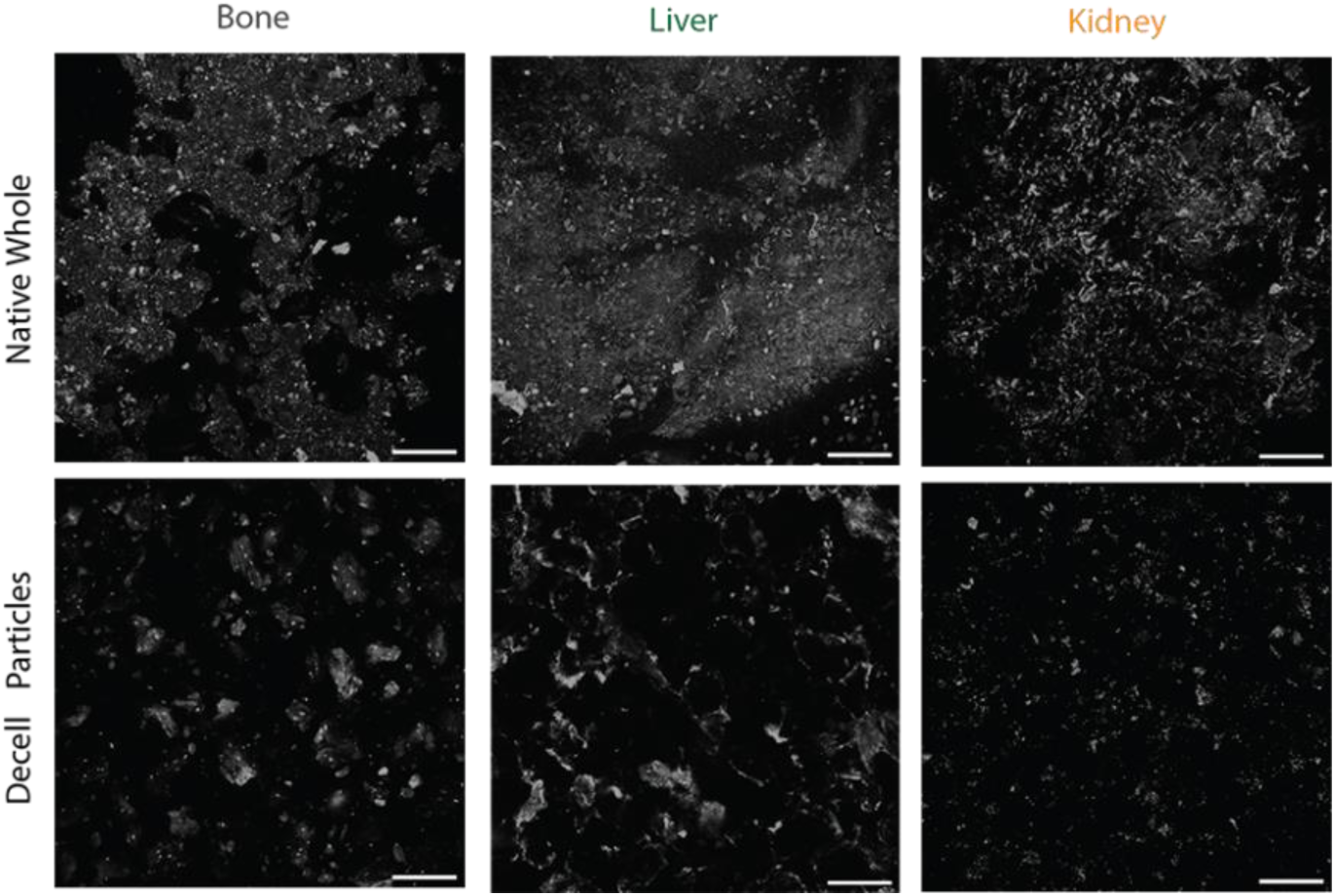
Confocal visualization of DNA content for bone, liver, and kidney. Representative confocal images of bone, liver, and kidney tissue showing DAPI-stained nuclei (white) in native whole tissue and decell ECM particles. Scale bars = 200 µm.

**Supplemental Figure 2.**
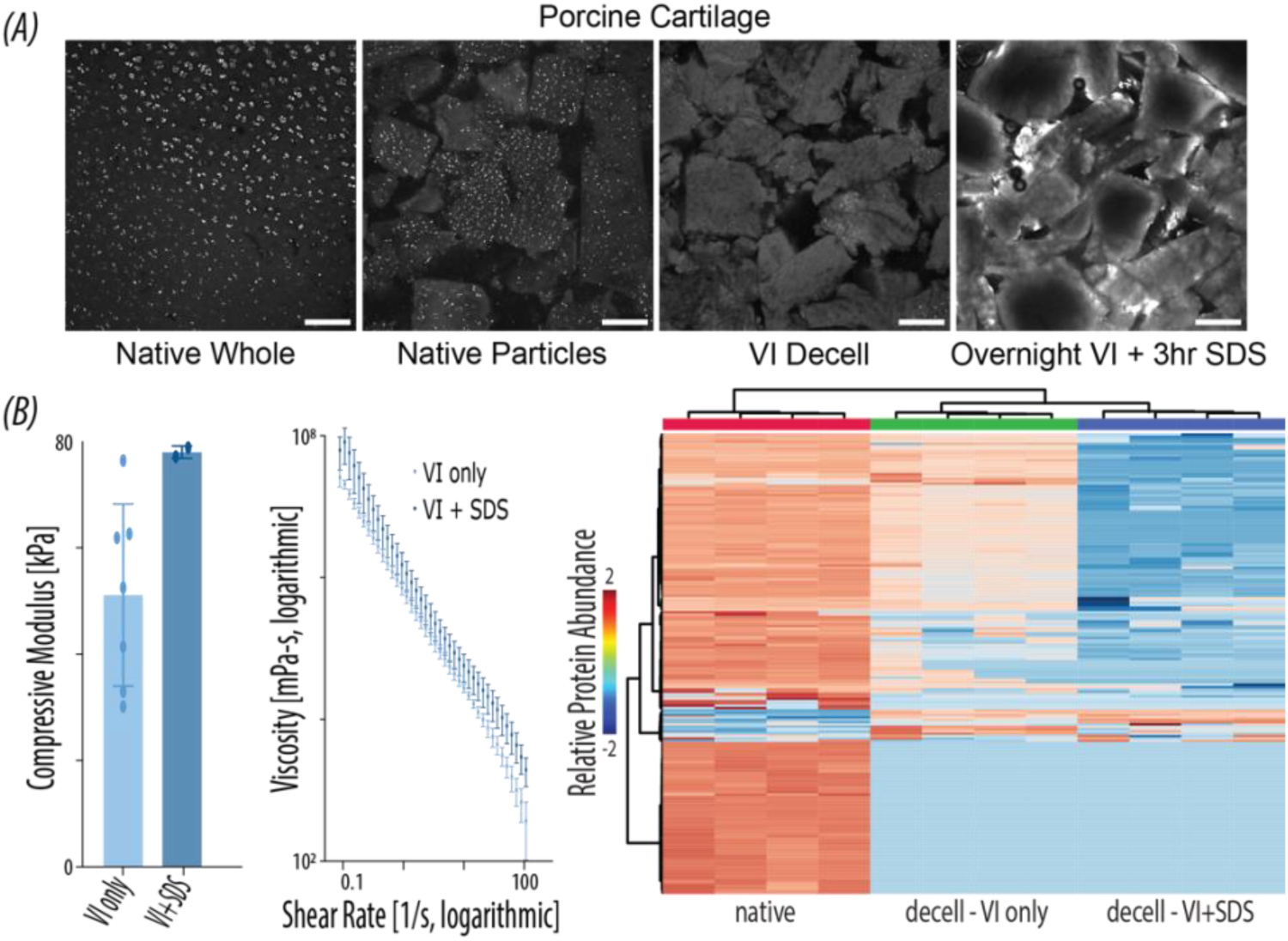
Comparison between cartilage processed with viral inactivation-only versus overnight viral inactivation plus detergent treatment. (A) Confocal images of cartilage tissue showing DAPI-stained nuclei (white) in native whole tissue, native ECM particles, viral-inactivation-only (VI only) decell particles, and overnight viral inactivation plus detergent (VI+SDS) decell particles (scale bars = 200µm). (B) Viscosity (N = 3-10) (left) and compressive modulus (N = 2-7) (center) of VI-only vs. VI + SDS cartilage gECM. Heatmap displays relative protein abundance of Native cartilage, Decell VI-only cartilage, and Decell VI + SDS cartilage particles.

**Supplemental Figure 3.**
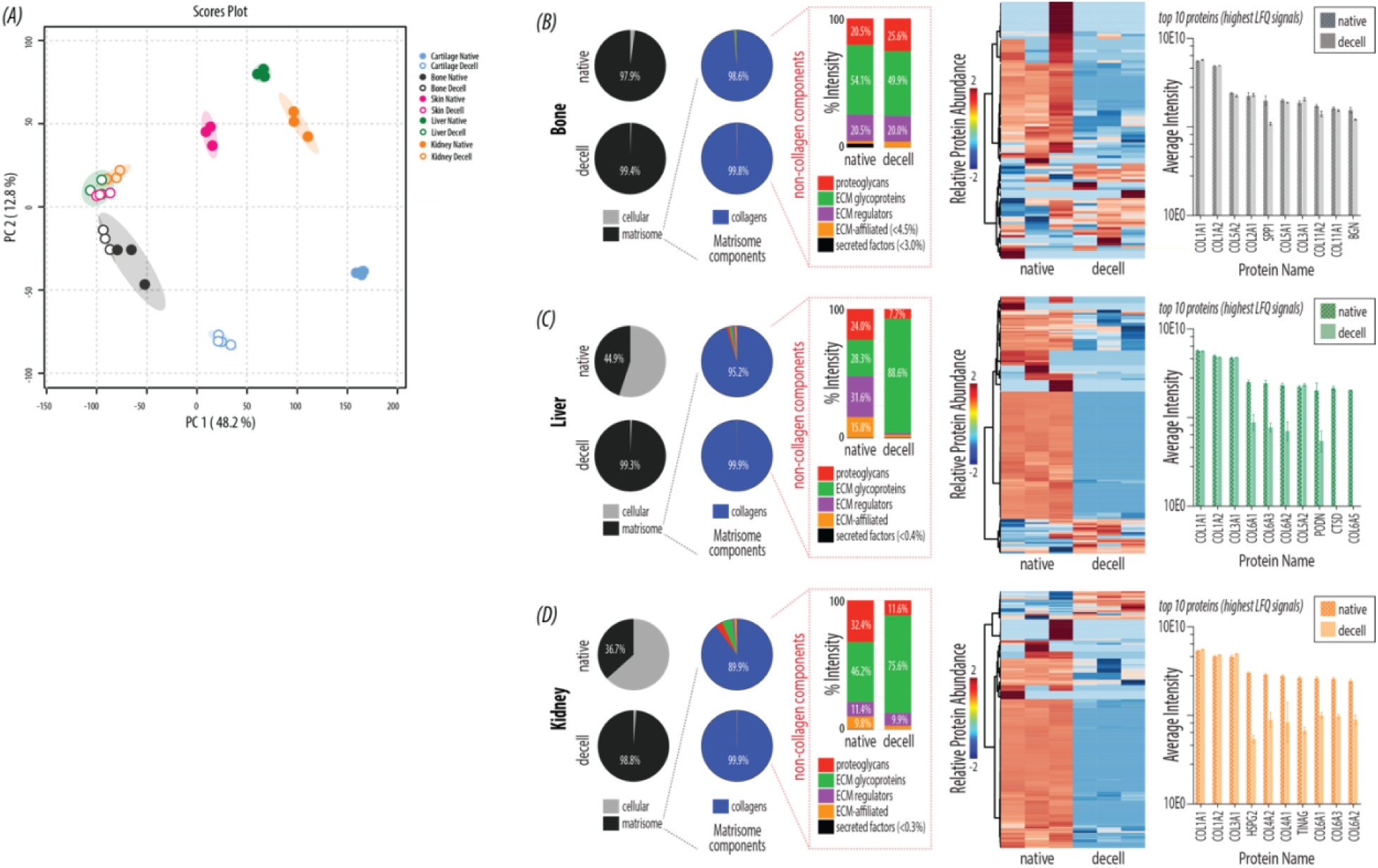
Proteomics for all tissues. (A) PCA plot for native and decell particles (N = 3) for all tissues (cartilage, bone, skin, liver, kidney). (B-D) Proteomic analysis (N = 3) of native vs. decell bone (B), liver (C), and kidney (D) – For each tissue, we present cellular vs. matrisomal protein content by LFQ signal. Remaining plots portray matrisome proteins only – matrisome subcategory distribution by LFQ signal, heat map with relative protein abundance, and top 10 protein abundance. Error bars = standard deviation.

**Supplemental Figure 4.**
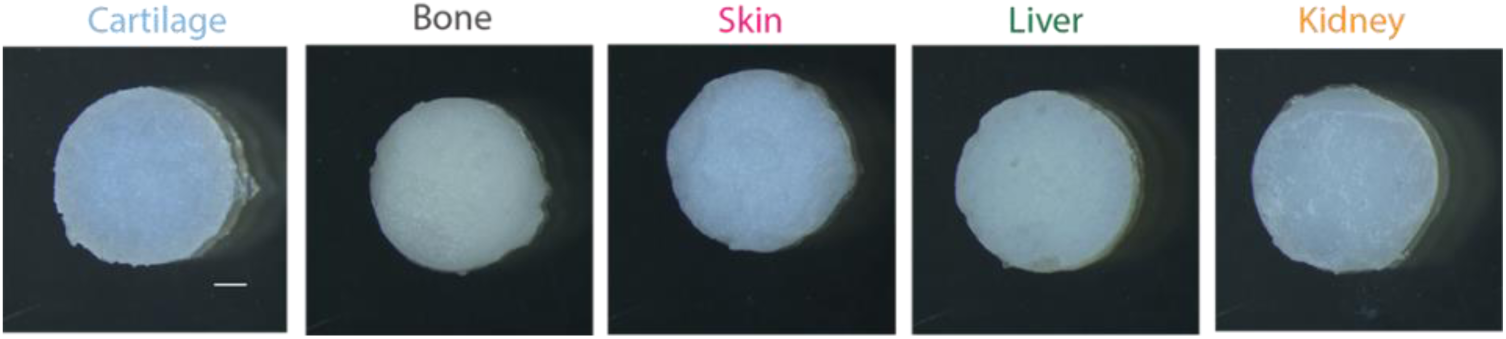
Stereoscopic images of polymerized gECM. Top-down images of polymerized gECM constructs for all tissues. Scale bar = 1 mm.

**Supplemental Figure 5.**
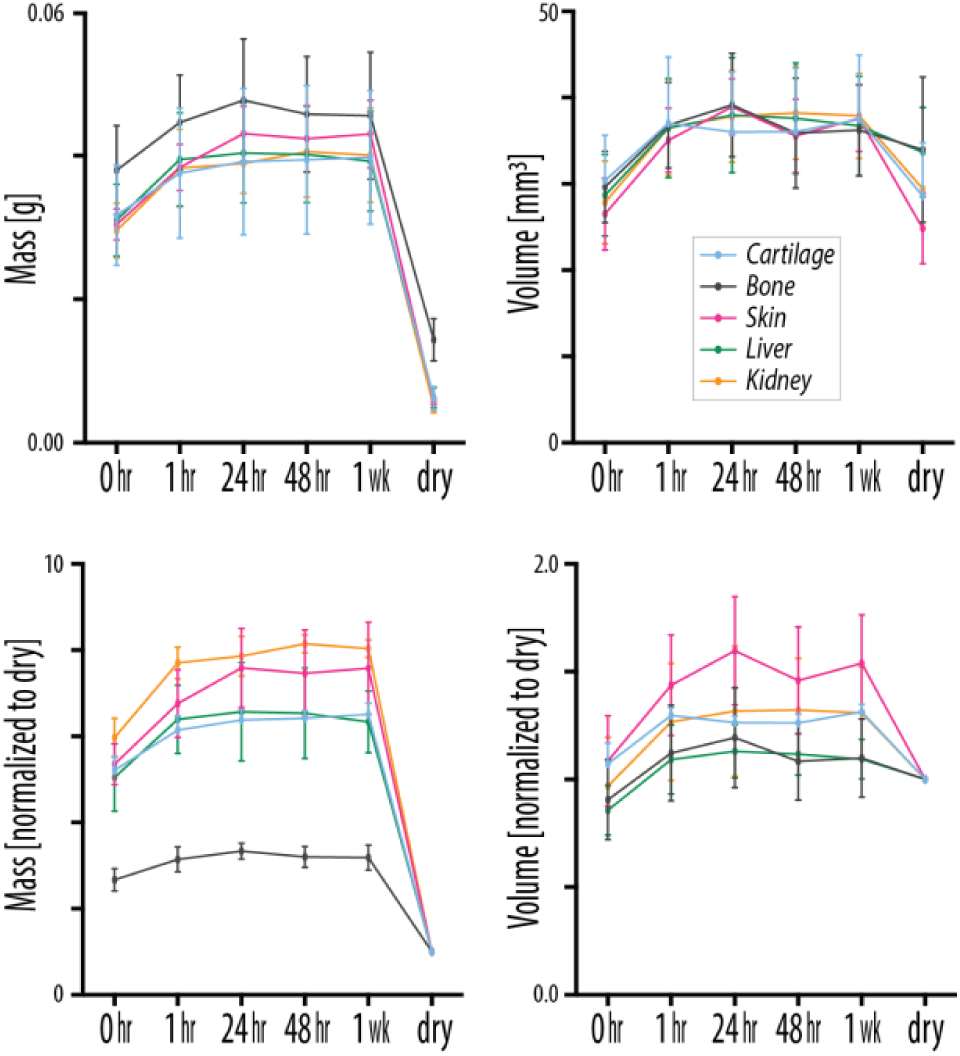
Swelling data raw values and normalized to dry for gECM hydrogels. Mass and volume for all tissue gECM hydrogels over 1 week swelling in DPBS. Shown here as raw values, and values normalized to the dry (0hr) mass or volume.

**Supplemental Figure 6.**
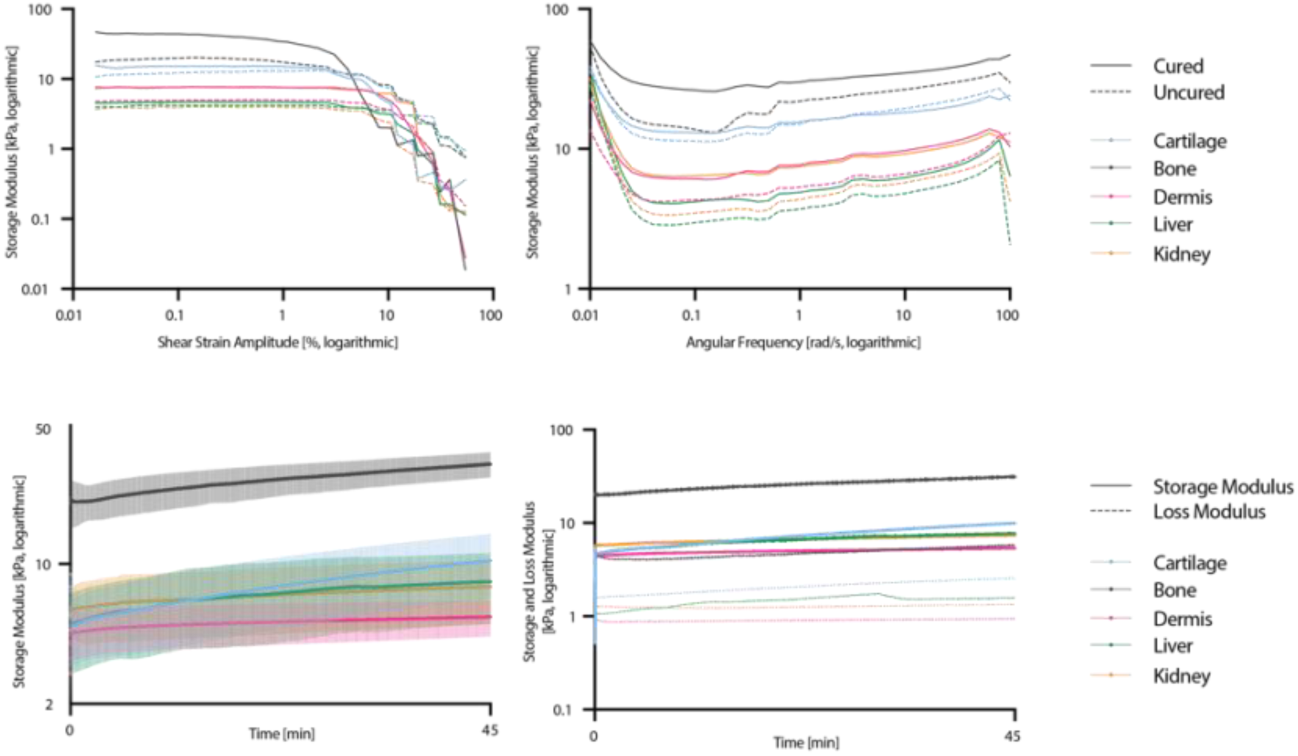
Rheological amplitude, frequency, and time sweeps. Amplitude and frequency sweeps for all tissue gECM hydrogels in a noncured (immediately after mixing) and cured (polymerized 45 mins at 37°C) state (N = 1). Raw storage modulus over a 45-minute time sweep (N = 4) (error bars = standard deviation). Raw mean storage and loss modulus over a 45-minute time sweep (N = 4).

**Supplemental Figure 7.**
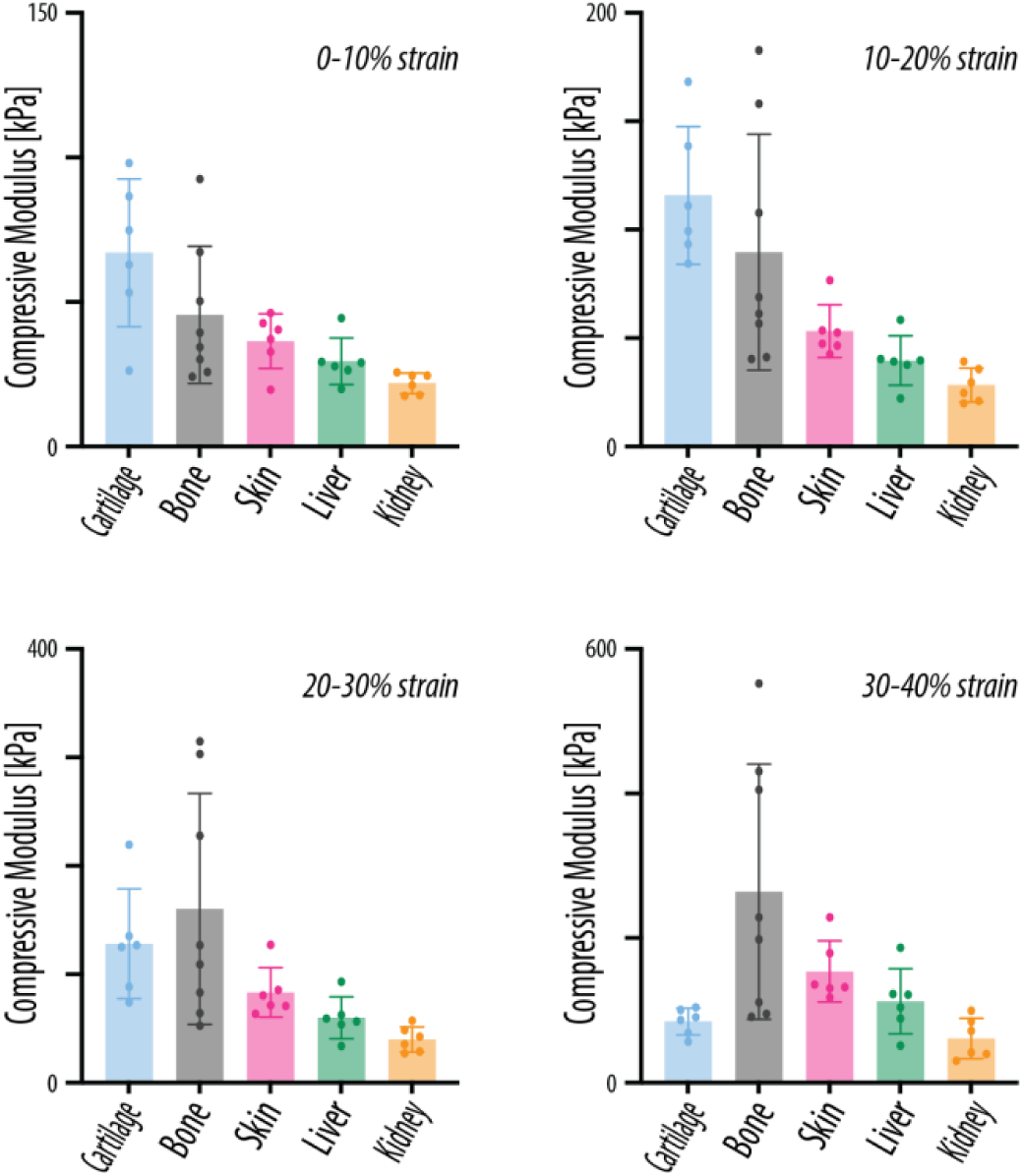
Compressive modulus calculated for various strain windows. Compressive modulus calculated at 0-10%, 10-20%, 20-30%, and 30-40% strain for all tissue gECM hydrogels (N = 6-8). Error bars = standard deviation.

## References

1. [1] 3D Bioprinting Market to Set Astonishing Growth (CAGR of 21.3%) n.d. https://www.openpr.com/news/1721502/3d-bioprinting-market-to-set-astonishing-growth-cagr-of-21-3-by-key-players-biobots-cellink-gesim-poietis.html (accessed December 29, 2025).

[2] Barthold JE, McCreery KP, Martinez J, Bellerjeau C, Ding Y, Bryant SJ, et al. Particulate ECM biomaterial ink is 3D printed and naturally crosslinked to form structurally-layered and lubricated cartilage tissue mimics. Biofabrication 2022;14:025021. 10.1088/1758-5090/AC584C.

[3] Aguilera SB, McCarthy A, Khalifian S, Lorenc ZP, Goldie K, Chernoff WG. The Role of Calcium Hydroxylapatite (Radiesse) as a Regenerative Aesthetic Treatment: A Narrative Review. Aesthet Surg J 2023;43:1063–90. 10.1093/ASJ/SJAD173.

[4] Murphy SV, Atala A. Organ engineering - combining stem cells, biomaterials, and bioreactors to produce bioengineered organs for transplantation. BioEssays 2013;35:163–72. 10.1002/BIES.201200062.

5. [5] Medical Devices Containing Materials Derived from Animal Sources (Except for In Vitro Diagnostic Devices) Guidance for Industry and Food and Drug Administration Staff Preface Public Comment n.d.

6. [6] Fda, Cber. Considerations for the Use of Human-and Animal-Derived Materials in the Manufacture of Cellular and Gene Therapy and Tissue-Engineered Medical Products Draft Guidance for Industry n.d.

[7] Zushin PJH, Mukherjee S, Wu JC. FDA Modernization Act 2.0: transitioning beyond animal models with human cells, organoids, and AI/ML-based approaches. J Clin Invest 2023;133:e175824. 10.1172/JCI175824.

[8] Han JJ. FDA Modernization Act 2.0 allows for alternatives to animal testing. Artif Organs 2023;47:449–50. 10.1111/AOR.14503.

[9] Liu X, Yue T, Kojima M, Huang Q, Arai T. Bio-assembling and Bioprinting for Engineering Microvessels from the Bottom Up. Int J Bioprint 2021;7:3–17. 10.18063/IJB.V7I3.366.

[10] Kim PH, Yim HG, Choi YJ, Kang BJ, Kim J, Kwon SM, et al. Injectable multifunctional microgel encapsulating outgrowth endothelial cells and growth factors for enhanced neovascularization. J Control Release 2014;187:1–13. 10.1016/J.JCONREL.2014.05.010.

[11] Siltanen C, Yaghoobi M, Haque A, You J, Lowen J, Soleimani M, et al. Microfluidic fabrication of bioactive microgels for rapid formation and enhanced differentiation of stem cell spheroids. Acta Biomater 2016;34:125–32. 10.1016/J.ACTBIO.2016.01.012.

[12] Chu H, Zhang K, Rao Z, Song P, Lin Z, Zhou J, et al. Harnessing decellularised extracellular matrix microgels into modular bioinks for extrusion-based bioprinting with good printability and high post-printing cell viability. Biomaterials Translational 2023;4:115. 10.12336/biomatertransl.2023.02.006.

[13] Luo SC, Li MT, Wang YC, Kankala RK, Wang S Bin, Xu PY, et al. Microfluidic GelMA/Bone-Derived Extracellular Matrix Microgels for Enhanced Stem Cell-Based Bone Regeneration. ACS Appl Bio Mater 2026;9:906–20. 10.1021/acsabm.5c01821.

[14] Daly AC, Riley L, Segura T, Burdick JA. Hydrogel microparticles for biomedical applications. Nature Reviews Materials 2019 5:1 2019;5:20–43. 10.1038/s41578-019-0148-6.

[15] Rowley AT, Nagalla RR, Wang SW, Liu WF. Extracellular Matrix-Based Strategies for Immunomodulatory Biomaterials Engineering. Adv Healthc Mater 2019;8. 10.1002/ADHM.201801578.

[16] Cramer MC, Badylak SF. Extracellular Matrix-Based Biomaterials and Their Influence Upon Cell Behavior. Ann Biomed Eng 2020;48:2132. 10.1007/S10439-019-02408-9.

[17] Novak T, Seelbinder B, Twitchell CM, Voytik-Harbin SL, Neu CP. Dissociated and Reconstituted Cartilage Microparticles in Densified Collagen Induce Local hMSC Differentiation. Adv Funct Mater 2016;26:5427. 10.1002/ADFM.201601877.

[18] Barthold JE, St Martin BM, Lalitha Sridhar S, Vernerey F, Ellyse Schneider S, Wacquez A, et al. Recellularization and Integration of Dense Extracellular Matrix by Percolation of Tissue Microparticles. Adv Funct Mater 2021;31:2103355. 10.1002/ADFM.202103355.

[19] Pati F, Jang J, Ha DH, Won Kim S, Rhie JW, Shim JH, et al. Printing three-dimensional tissue analogues with decellularized extracellular matrix bioink. Nature Communications 2014 5:1 2014;5:1–11. 10.1038/ncomms4935.

[20] Ahn G, Min KH, Kim C, Lee JS, Kang D, Won JY, et al. Precise stacking of decellularized extracellular matrix based 3D cell-laden constructs by a 3D cell printing system equipped with heating modules. Sci Rep 2017;7. 10.1038/S41598-017-09201-5.

[21] Neto MD, Oliveira MB, Mano JF. Microparticles in Contact with Cells: From Carriers to Multifunctional Tissue Modulators. Trends Biotechnol 2019;37:1011–28. 10.1016/J.TIBTECH.2019.02.008.

[22] Townsend JM, Dennis SC, Whitlow J, Feng Y, Wang J, Andrews B, et al. Colloidal Gels with Extracellular Matrix Particles and Growth Factors for Bone Regeneration in Critical Size Rat Calvarial Defects. AAPS J 2017;19:703–11. 10.1208/S12248-017-0045-0.

[23] Fedorovich NE, Wijnberg HM, Dhert WJA, Alblas J. Distinct tissue formation by heterogeneous printing of osteo-and endothelial progenitor cells. Tissue Eng Part A 2011;17:2113–21. 10.1089/TEN.TEA.2011.0019/ASSET/IMAGES/LARGE/FIGURE5.JPEG.

[24] Kim MK, Jeong W, Lee SM, Kim JB, Jin S, Kang HW. Decellularized extracellular matrix-based bio-ink with enhanced 3D printability and mechanical properties. Biofabrication 2020;12. 10.1088/1758-5090/AB5D80.

[25] Isaeva E V., Beketov EE, Demyashkin GA, Yakovleva ND, Arguchinskaya N V., Kisel AA, et al. Cartilage Formation In Vivo Using High Concentration Collagen-Based Bioink with MSC and Decellularized ECM Granules. Int J Mol Sci 2022;23. 10.3390/IJMS23052703.

[26] Jung CS, Kim BK, Lee J, Min BH, Park SH. Development of Printable Natural Cartilage Matrix Bioink for 3D Printing of Irregular Tissue Shape. Tissue Eng Regen Med 2018;15:155. 10.1007/S13770-017-0104-8.

[27] Kara A, Distler T, Polley C, Schneidereit D, Seitz H, Friedrich O, et al. 3D printed gelatin/decellularized bone composite scaffolds for bone tissue engineering: Fabrication, characterization and cytocompatibility study. Mater Today Bio 2022;15. 10.1016/J.MTBIO.2022.100309.

[28] Gharacheh H, Guvendiren M. Cell-Laden Composite Hydrogel Bioinks with Human Bone Allograft Particles to Enhance Stem Cell Osteogenesis. Polymers (Basel) 2022;14:3788. 10.3390/POLYM14183788.

[29] Ratheesh G, Vaquette C, Xiao Y. Patient-Specific Bone Particles Bioprinting for Bone Tissue Engineering. Adv Healthc Mater 2020;9. 10.1002/ADHM.202001323.

[30] Hung BP, Naved BA, Nyberg EL, Dias M, Holmes CA, Elisseeff JH, et al. Three-Dimensional Printing of Bone Extracellular Matrix for Craniofacial Regeneration. ACS Biomater Sci Eng 2016;2:1806–16. 10.1021/ACSBIOMATERIALS.6B00101.

[31] Thitiset T, Damrongsakkul S, Bunaprasert T, Leeanansaksiri W, Honsawek S. Development of collagen/demineralized bone powder scaffolds and periosteum-derived cells for bone tissue engineering application. Int J Mol Sci 2013;14:2056–71. 10.3390/IJMS14012056.

[32] Ghorbani F, Ekhtiari M, Moeini Chaghervand B, Moradi L, Mohammadi B, Kajbafzadeh AM. Detection of the residual concentration of sodium dodecyl sulfate in the decellularized whole rabbit kidney extracellular matrix. Cell Tissue Bank 2022;23:119–28. 10.1007/S10561-021-09921-Z.

[33] Cameron R, Smith K. Virus clearance methods applied in bioprocessing operations: an overview of selected inactivation and removal methods. Pharm Bioprocess 2014;2:75–83. 10.4155/PBP.13.61.

[34] Crapo PM, Gilbert TW, Badylak SF. An overview of tissue and whole organ decellularization processes. Biomaterials 2011;32:3233–43. 10.1016/J.BIOMATERIALS.2011.01.057.

[35] Alcaide-Ruggiero L, Cugat R, Domínguez JM. Proteoglycans in Articular Cartilage and Their Contribution to Chondral Injury and Repair Mechanisms. Int J Mol Sci 2023;24:10824. 10.3390/ijms241310824.

[36] Smith MM, Melrose J. Proteoglycans in Normal and Healing Skin. Adv Wound Care (New Rochelle) 2015;4:152. 10.1089/wound.2013.0464.

[37] Scott AK, Gallagher KM, Schneider SE, Kurse A, Neu CP. Epigenetic Priming Enhances Chondrogenic Potential of Expanded Chondrocytes. Tissue Eng Part A 2024;30:415–25. 10.1089/TEN.TEA.2023.0170.

[38] Scott AK, Casas E, Schneider SE, Swearingen AR, Van Den Elzen CL, Seelbinder B, et al. Mechanical memory stored through epigenetic remodeling reduces cell therapeutic potential. Biophys J 2023;122:1428–44. 10.1016/j.bpj.2023.03.004.

[39] Qazi TH, Muir VG, Burdick JA. Methods to Characterize Granular Hydrogel Rheological Properties, Porosity, and Cell Invasion. ACS Biomater Sci Eng 2022;8:1427–42. 10.1021/ACSBIOMATERIALS.1C01440.

[40] Kan X, Zhang S, Kwok E, Chu Y, Chen L, Zeng X. Granular hydrogels with tunable properties prepared from gum Arabic and protein microgels. Int J Biol Macromol 2024;273:132878. 10.1016/J.IJBIOMAC.2024.132878.

[41] Feng S, Chen K, Wang S. Practical Guide to the Design of Granular Hydrogels for Customizing Complex Cellular Microenvironments. Adv Healthc Mater 2025;14:e01947. 10.1002/ADHM.202501947.

[42] Deptuła P, Łysik D, Pogoda K, Cieśluk M, Namiot A, Mystkowska J, et al. Tissue Rheology as a Possible Complementary Procedure to Advance Histological Diagnosis of Colon Cancer. ACS Biomater Sci Eng 2020;6:5620–31. 10.1021/ACSBIOMATERIALS.0C00975.

[43] Emiroglu DB, Bekcic A, Dranseikiene D, Zhang X, Zambelli T, deMello AJ, et al. Building block properties govern granular hydrogel mechanics through contact deformations. Sci Adv 2022;8. 10.1126/sciadv.add8570.

[44] Muir VG, Qazi TH, Shan J, Groll J, Burdick JA. Influence of Microgel Fabrication Technique on Granular Hydrogel Properties. ACS Biomater Sci Eng 2021;7:4269–81. 10.1021/ACSBIOMATERIALS.0C01612.

[45] Muir VG, Weintraub S, Dhand AP, Fallahi H, Han L, Burdick JA. Influence of Microgel and Interstitial Matrix Compositions on Granular Hydrogel Composite Properties. Advanced Science 2023;10:2206117. 10.1002/advs.202206117DigitalObjectIdentifier(DOI).

[46] Si L, Bai H, Rodas M, Cao W, Oh CY, Jiang A, et al. A human-airway-on-a-chip for the rapid identification of candidate antiviral therapeutics and prophylactics. Nature Biomedical Engineering 2021 5:8 2021;5:815–29. 10.1038/s41551-021-00718-9.

[47] Ghosh S, Seelbinder B, Henderson JT, Watts RD, Scott AK, Veress AI, et al. Deformation Microscopy for Dynamic Intracellular and Intranuclear Mapping of Mechanics with High Spatiotemporal Resolution. Cell Rep 2019;27:1607–1620.e4. 10.1016/J.CELREP.2019.04.009.

[48] Guo Y, Song Y, Xiong S, Wang T, Liu W, Yu Z, et al. Mechanical Stretch Induced Skin Regeneration: Molecular and Cellular Mechanism in Skin Soft Tissue Expansion. Int J Mol Sci 2022;23:9622. 10.3390/ijms23179622.

[49] Liang X, Huang X, Zhou Y, Jin R, Li Q. Mechanical Stretching Promotes Skin Tissue Regeneration via Enhancing Mesenchymal Stem Cell Homing and Transdifferentiation. Stem Cells Transl Med 2016;5:960–9. 10.5966/sctm.2015-0274.

[50] Occhetta P. Hyperphysiological compression of articular cartilage induces an osteoarthritic phenotype in a cartilage-on-a-chip model _ Enhanced Reader. Nature BME 2019;3:545–57.

[51] Chan DD, Cai L, Butz KD, Trippel SB, Nauman EA, Neu CP. In vivo articular cartilage deformation: noninvasive quantification of intratissue strain during joint contact in the human knee. Scientific Reports 2016 6:1 2016;6:19220-. 10.1038/srep19220.

[52] Ghosh S, Scott AK, Seelbinder B, Barthold JE, Martin BMS, Kaonis S, et al. Dedifferentiation alters chondrocyte nuclear mechanics during in vitro culture and expansion. Biophys J 2021;121:131. 10.1016/J.BPJ.2021.11.018.

[53] Chen Y, Yu Y, Wen Y, Chen J, Lin J, Sheng Z, et al. A high-resolution route map reveals distinct stages of chondrocyte dedifferentiation for cartilage regeneration. Bone Research 2022 10:1 2022;10:38-. 10.1038/s41413-022-00209-w.

[54] Ledoult E, Jendoubi M, Collet A, Guerrier T, Largy A, Speca S, et al. Simple gene signature to assess murine fibroblast polarization. Scientific Reports 2022 12:1 2022;12:11748-. 10.1038/s41598-022-15640-6.

[55] Łuszczyński K, Soszyńska M, Komorowski M, Lewandowska P, Zdanowski R, Sobiepanek A, et al. Markers of Dermal Fibroblast Subpopulations for Viable Cell Isolation via Cell Sorting: A Comprehensive Review. Cells 2024;13:1206. 10.3390/CELLS13141206.

[56] Simman R. Role of small intestinal submucosa extracellular matrix in advanced regenerative wound therapy. J Wound Care 2023;32:S3–10. 10.12968/JOWC.2023.32.SUP1A.S3.

[57] Sakata N, Yoshimatsu G, Kawakami R, Chinen K, Aoyagi C, Kodama S. Procedure of Adult Porcine Islet Isolation. Tissue Eng Part C Methods 2023;29:144–53. 10.1089/TEN.TEC.2023.0020.

[58] Hua K chi, Feng J tao, Yang X gang, Wang F, Zhang H, Yang L, et al. Assessment of the Defatting Efficacy of Mechanical and Chemical Treatment for Allograft Cancellous Bone and Its Effects on Biomechanics Properties of Bone. Orthop Surg 2020;12:617–30. 10.1111/OS.12639.

[59] Trefalt G, Cao T, Sugimoto T, Borkovec M. Heteroaggregation between Charged and Neutral Particles. Langmuir 2020;36:5303–11. 10.1021/ACS.LANGMUIR.0C00667.

[60] Zhang L, Haddouti EM, Welle K, Burger C, Wirtz DC, Schildberg FA, et al. The Effects of Biomaterial Implant Wear Debris on Osteoblasts. Front Cell Dev Biol 2020;8:525428. 10.3389/FCELL.2020.00352/FULL.

[61] Wang K, Wu J, Day RE, Kirk TB. Utilizing confocal microscopy to measure refractive index of articular cartilage. J Microsc 2012;248:281–91. 10.1111/j.1365-2818.2012.03674.x DigitalObjectIdentifier(DOI).

[62] Cardinali MA, Dallari D, Govoni M, Stagni C, Marmi F, Tschon M, et al. Brillouin micro-spectroscopy of subchondral, trabecular bone and articular cartilage of the human femoral head. Biomed Opt Express 2019;10:2606. 10.1364/BOE.10.002606.

[63] Knüttel A, Boehlau-Godau M. Spatially confined and temporally resolved refractive index and scattering evaluation in human skin performed with optical coherence tomography. J Biomed Opt 2000;5:83. 10.1117/1.429972.

[64] Giannios P, Toutouzas KG, Matiatou M, Stasinos K, Konstadoulakis MM, Zografos GC, et al. Visible to near-infrared refractive properties of freshly-excised human-liver tissues: marking hepatic malignancies. Scientific Reports 2016 6:1 2016;6:27910-. 10.1038/srep27910.

[65] Young PA, Clendenon SG, Byars JM, Dunn KW. The effects of refractive index heterogeneity within kidney tissue on multiphoton fluorescence excitation microscopy. J Microsc 2011;242:148–56. 10.1111/j.1365-2818.2010.03448.xDigitalObjectIdentifier(DOI).

[66] Ebert DW, Roberts CJ, Farrar SK, Johnston WM, Litsky AS, Bertone AL. Articular cartilage optical properties in the spectral range 300--850 nm. Https://DoiOrg/101117/1429893 1998;3:326–33. 10.1117/1.429893.

[67] Shimojo Y, Nishimura T, Hazama H, Ozawa T, Awazu K. Measurement of absorption and reduced scattering coefficients in Asian human epidermis, dermis, and subcutaneous fat tissues in the 400- to 1100-nm wavelength range for optical penetration depth and energy deposition analysis. J Biomed Opt 2020;25:1. 10.1117/1.JBO.25.4.045002.

[68] Botelho AR, Silva HF, Martins IS, Carneiro IC, Carvalho SD, Henrique RM, et al. Fast calculation of spectral optical properties and pigment content detection in human normal and pathological kidney. Spectrochim Acta A Mol Biomol Spectrosc 2023;286. 10.1016/j.saa.2022.122002.

[69] Bashkatov AN, Genina EA, Kochubey VI, Tuchin V V., Bashkatov AN, Genina EA, et al. Optical properties of human cranial bone in the spectral range from 800 to 2000 nm. SPIE 2006;6163:616310. 10.1117/12.697305.

[70] Wiśniewski JR, Zougman A, Nagaraj N, Mann M. Universal sample preparation method for proteome analysis. Nature Methods 2009 6:5 2009;6:359–62. 10.1038/nmeth.1322.

[71] Kong AT, Leprevost F V., Avtonomov DM, Mellacheruvu D, Nesvizhskii AI. MSFragger: ultrafast and comprehensive peptide identification in mass spectrometry-based proteomics. Nat Methods 2017;14:513–20. 10.1038/nmeth.4256.

[72] Mellacheruvu D, Wright Z, Couzens AL, Lambert JP, St-Denis NA, Li T, et al. The CRAPome: a contaminant repository for affinity purification–mass spectrometry data. Nature Methods 2013 10:8 2013;10:730–6. 10.1038/nmeth.2557.

[73] Shao X, Taha IN, Clauser KR, Gao Y (Tom), Naba A. MatrisomeDB: the ECM-protein knowledge database. Nucleic Acids Res 2020;48:D1136–44. 10.1093/nar/gkz849.

